# In situ generated vaccine-like pyroptosome for personalized cancer immunotherapy

**DOI:** 10.1101/2025.02.09.636745

**Authors:** Binlong Chen, Fangjie Wan, Heming Xia, Xingquan Pan, Letong Wang, Yaoqi Wang, Yue Yan, Jianxiong Liu, Mingmei Tang, Ye Yang, Mengmeng Qin, Jiaona Ren, Lijun Zhong, Wei Chen, Qiang Zhang, Yiguang Wang

## Abstract

The efficacy of in situ cancer vaccination has been hampered by poor spatiotemporal orchestration of multiple key steps of cancer-immunity cycle in most tumours and systemic toxicity related to therapeutic strategies. Here, we report a systemic injectable and pyroptosis-enabled nanoadjuvant (SPEN) that precisely evokes the secretion of vaccine-like pyroptosome in tumour area for eliciting robust anti-tumour immunity. SPEN induces vigorous immunogenic pyroptosis, triggering the efficient release of tumour antigen-rich pyroptosomes, DAMPs, and proinflammatory cytokines. Upon the activatable release of a TLR7/8 agonist into pyroptosome, the generated pyroptosome functions as in situ cancer vaccine for cooperatively activating the cancer-immunity cycle while avoiding systemic toxicity. This in situ vaccine boosts both innate and adaptive immune response, facilitating the eradication of primary tumour and long-lasting cancer prevention. Our findings provide new insights into the rational design of pyroptosis-inducing nanomedicines for boosting the cancer-immunity cycle, thus advancing personalized cancer immunotherapy.

## Introduction

Cancer vaccines, as a leading strategy in immunotherapies, hold great potential to boost the adaptive immune response and enhance anti-tumour immunity. Although numerous contributions have been made in pre-clinical and clinical investigation of cancer vaccines over the past decades, they have yet to become a hallmark therapy in oncology^1^. The outcome of vaccines largely depends on their immunogenicity, which includes antigen and adjuvant. However, because of inter-tumour and inter-patient heterogeneity of tumour antigens, there is an urgent need to develop personalized neoantigen vaccine^2, 3^. Benefiting from the emergence of genome sequencing, machine learning, and mRNA technology, personalized vaccine therapy has achieved promising outcomes in pre-clinical and clinical studies^4, 5^. Nevertheless, the high cost and complex manufacturing challenge the generalization of the customized patient-specific cancer vaccine^6^.

In situ cancer vaccination requires the orchestrated activation of cancer-immunity cycles (CIC), including the antigen release from dying cancer cells, antigen processing and presentation by antigen-presenting cells (APCs), the generation of tumour-specific effector T cells for lysis of tumour cells, which further boosts the CIC^7, 8^. However, the rate-limiting capability of each successive step in the CIC and the immunosuppressive tumour microenvironments pose significant challenges to the efficient activation of CIC and subsequent anti-tumour immune response^9^. Numerous strategies have been developed to amplify CIC, including hybridization with bacterial membranes^10^, cooperation with fluoropolymer^11^, combination with immune adjuvants^12^, integration with mRNA therapy^13^, and the extensively investigated chemo/radiotherapy-mediated immunogenic cell death (ICD)^14^. Accumulating evidence has revealed that the death nature dictates the initiation of CIC^15, 16^. Apoptosis has been recently recognized as a largely immune-silent form of cell death, characterized by self-degradation and minimal release of cellular content^17^. In contrast, the lytic nature of programed necrosis optimizes the provision of antigens and adjuvanticity, thereby boosting CIC for a potent tumour-specific T cell immunity. Thus, there is an urgent need to exploit necroptotic cell death with efficient cascade amplification of CIC for in situ cancer vaccination.

Recently, we have developed a pyroptosis nanotuner technology that spatiotemporally evokes distinct endocytic organelle stress during endosome maturation for precisely regulating pyroptosis and apoptosis of cancer cells^18, 19^. Herein, we harness the nanotuner platform to engineer an in situ vaccine strategy through spatiotemporally fine-tuning ICD of cancer cells, boosting antigen presentation, and reprogramming the tumour microenvironment to close the cancer-immunity cycle (**Fig. 1**). The nanophotosensitizers with high membrane affinity efficiently amplify the immunogenic pyroptosis by early endosome (EE) stress, while negligible immunogenicity is provoked by lysosome (Ly) stress-mediated apoptosis. We further combine the ICD-tuning nanostrategy with oxidative stress-activatable TLR7/8 agonist to engineer a systemic injectable and pyroptosis-enabled nanoadjuvant (SPEN) for in situ generation of autologous cancer vaccine. After provoking pyroptosis-mediated eradication of primary tumours, SPEN generates pyroptosomes, which co-deliver tumour antigens and adjuvants into various immune cells. This pyroptosome-enabled vaccine boosts the anti-tumour innate and adaptive immune microenvironment within lymph nodes and tumours, facilitating up to 100% tumour eradication and 150-day long-lasting cancer prevention.

**Fig. 1.**
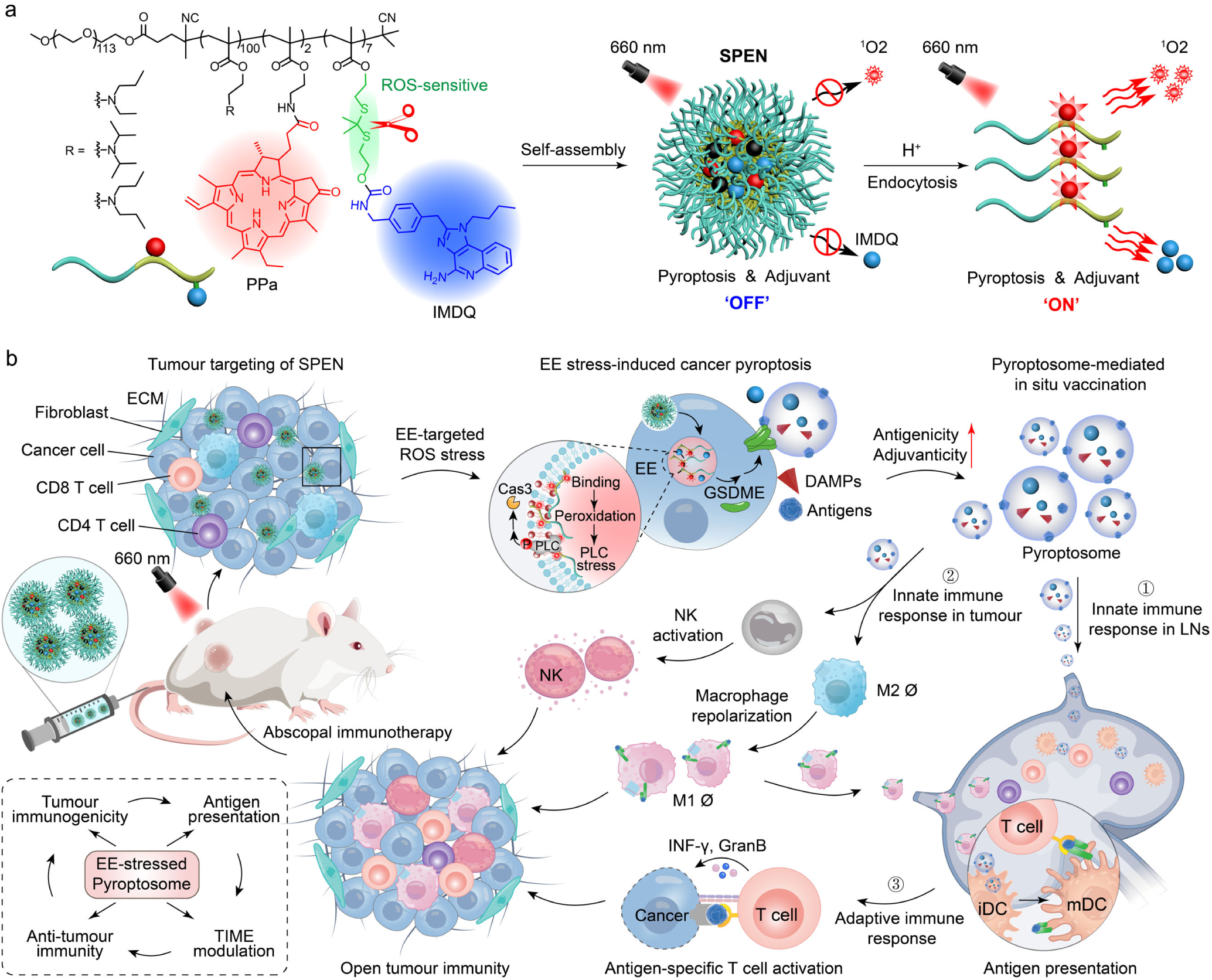
Schematic illustrations of the Systemic injectable and Pyroptosis-Enabled Nanoadjuvant (SPEN) for boosting cancer-immunity cycles (CIC) via in situ photogeneration of vaccine-like pyroptosome. **(a)** Design of the pH- and light-gated nanoadjuvant that elicits specific immunogenic pyroptosis of cancer cells via early endosome-targeted stress for in situ cancer vaccination. **(b)** SPEN keeps pyroptosis-inducing and adjuvant activities ‘OFF’ during blood circulation and extracellular distribution. After tumour accumulation and cellular endocytosis, SPEN specifically provokes cancer cell pyroptosis and release of toll-like receptor agonist, IMDQ, via early endosome-localized oxidative stress (SPEN_EE_), to activate the CIC and adjuvant activity simultaneously. Enabled by the highly efficient pyroptosis-inducing activity, SPEN_EE_ technology eradicates the primary tumours followed by the generation of vaccine-like pyroptosome, containing tumour antigens and IMDQ. By specifically amplifying the antigen presentation of macrophages and dendritic cells in tumour tissues, the pyroptosome functions as an in situ vaccine to prime the immune microenvironment of both the tumours and lymph nodes for robust and long-lasting cancer immunotherapy.

## Results

### Membrane affinity of nanophotosensitizer drives EE stress-induced cell pyroptosis

The nano-bio interaction is pivotal to the intracellular fate and theranostic outcomes of nanomedicine^20^. Based on our developed ultra-pH-sensitive nanotechnology, we first synthesized a series of amphiphilic copolymers mPEG_5k_-*b*-P(EPA_x_-*r*-DPA_100-x_), where x indicates the repeating units of 2-(ethylpropylamino) ethyl methacrylate (EPA) monomer in the total number of EPA and 2-(dipropylamino) ethyl methacrylate (DPA) monomers. The p*K*_a_ of synthetic copolymers ranged from 6.1 to 6.8, which falls within the pH range of early endosomes of cancer cells. Then, a small library of pyropheophorbide-a (PPa)-conjugated copolymers (**Supplementary Fig. 1**) was fabricated to explore the impacts of the chemical structures on the nano-bio interaction and thereafter pyroptosis-inducing activity of EE-targeted nanophotosensitizers (**Fig. 2a**). As shown in **Fig. 2b** and **Extended Fig. 1a,b**, the units of EPA monomers exhibited positive correlations with the cell membrane binding (R = 0.97) and membrane rupture-induced LDH release (R = 0.83) capacities of the ionizable nanophotosensitizers at pH 6.0. Moreover, introducing EPA monomers dramatically augmented the haemolytic efficacy of the protonated nanophotosensitizer-mediated oxidative stress (**Fig. 2c** and **Extended Fig. 1c**). The nanoprobes with distinct p*K*_a_^21^ exhibited tunable cellular distribution^18, 22^ and biological functions^23, 24^. To exclude the p*K*_a_ cue, we next synthesized PPa-conjugated PEG_5k_-*b*-poly(2-(diisopropylamino) ethyl methacrylate) (PEG_5k_-*b*-PDPA_100_) copolymer (abbreviated as PDPA) with the same p*K*_a_ of 6.5 and sharp pH response (ΔpH_on/off_ = 0.2) as PPa-conjugated mPEG_5k_-*b*-P(EPA_50_-*r*-DPA_50_) (abbreviated as PEPA) for parallel comparison (**Fig. 2d**). With the similar subcellular trafficking behaviour (**Extended Fig. 1d**), PEPA achieved 5.4-fold higher membrane binding (**Fig. 2e** and **Supplementary Fig. 2**) and 4.9-fold higher oxidative stress-induced haemolysis at pH 6.0 (**Extended Fig. 1e,f**) than PDPA. Then, we prepared the negatively-charged artificial lipid membranes (ALMs)^25^ to simulate the affinity between nanophotosensitizers and endo-lysosome membranes. Isothermal titration calorimetry (ITC) studies manifested that PEPA displayed potent binding affinity with the negatively charged ALMs at pH 6.0 rather than pH 7.4 (**Fig. 2f,g**). In contrast, PDPA exhibited negligible interaction with ALMs at pH 6.0 and pH 7.4 (**Supplementary Fig. 3**). Moreover, PPa was replaced by an environment-sensitive probe (Nile red)^26^ to report the interaction between ionizable copolymers and biomembranes. Results showed that PEPA exhibited a higher efficient binding with negative-charged ALMs than PDPA, while no significant difference was observed in their interaction with neutral ALMs (**Extended Fig. 1g**). The membrane-binding effect of nanophotosensitizer was also visualized by super-resolution microscopy. As shown in **Fig. 2h**, the doughnut-shaped fluorescence signals evidenced that PEPA presented a high binding affinity with endocytic organelle membranes over PDPA. The superior endosomal membrane affinity dramatically improved the cell photo-killing efficiency of PEPA when maturated into EE (PEPA_EE_). The half-maximal inhibitory concentration (IC_50_) of PDPA_EE_ was 3.5-fold higher than that of PEPA_EE_ (**Fig. 2i** and **Extended Fig. 1h**). The heightened cellular caspase-3 activation, oxidative stress signalling, cleavage of GSDME, and cell death phenotype studies collectively corroborated that PEPA_EE_ performed highly efficient and specific pyroptosis-inducing capacity (**Fig. 2j** and **Extended Fig. 1i-l**). These results demonstrated that intensifying the affinity of endosomal membrane augmented the nanophotosensitizer-mediated cell pyroptosis.

**Fig. 2.**
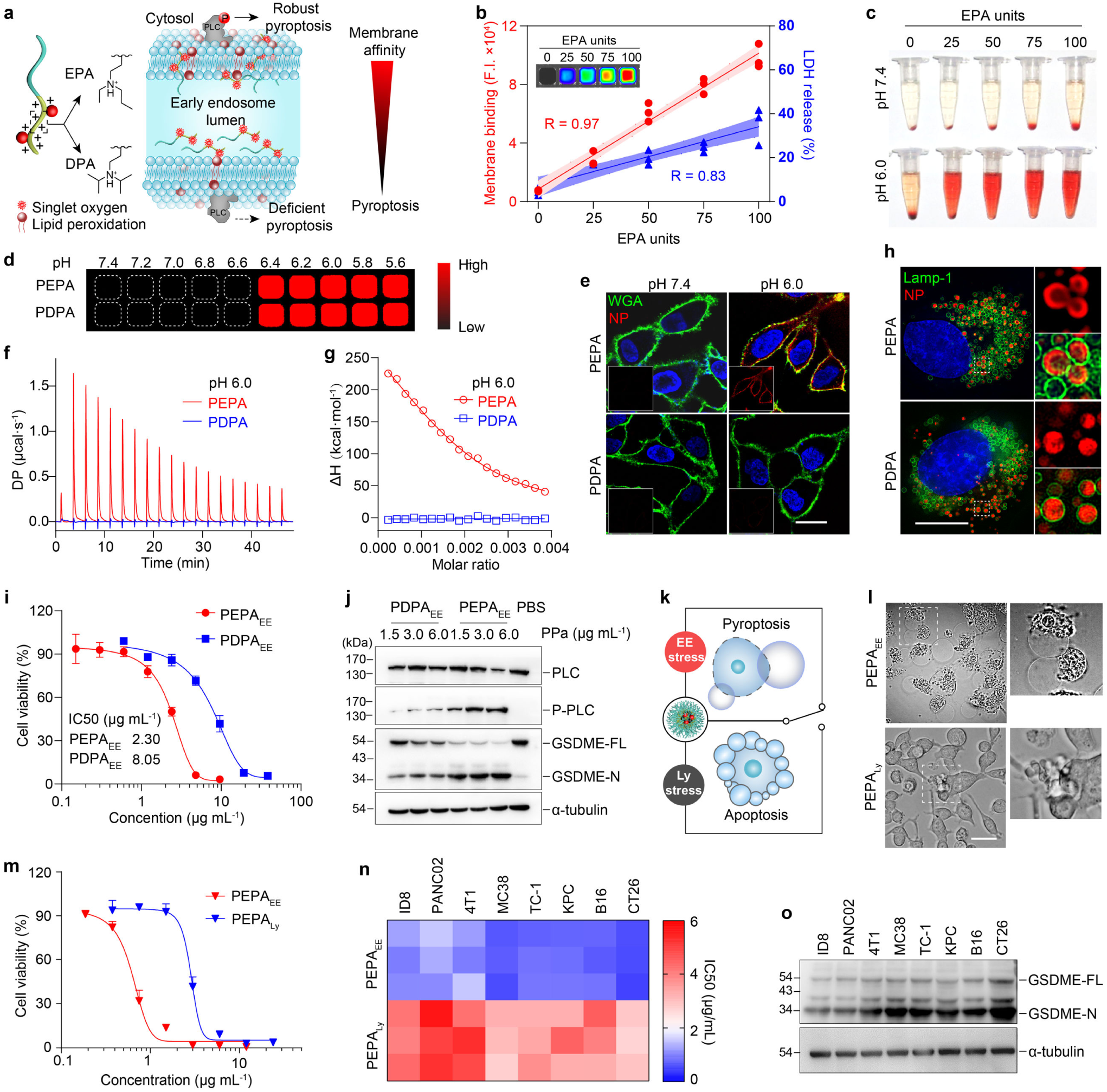
Membrane affinity of nanoparticle tunes EE stress-induced pyroptosis. **(a)** Schematic of the increased early endosomal membrane affinity of photosensitizer-functionalized ionizable copolymers for heightened pyroptosis-inducing activity. **(b)** The percentage of EPA monomers exhibits positive correlation with the A549 cell membrane binding ability (measured by flow cytometry and fluorescence imaging) and oxidative stress-induced LDH release of nanophotosensitizers at pH 6.0. The error band in dashed lines and shading shows the 95% confidence intervals of the fitted line by two-sided Student’s t-test analysis. **(c)** The membrane-binding of EPA-containing nanophotosensitizers evokes erythrocytes hemolysis at pH 6.0 rather than pH 7.4 under 660 nm irradiation (6 J cm^−2^). **(d)** Fluorescence images of PEPA and PDPA diluted in a series of PBS buffers with 0.2 pH unit increment in 384-well plate. **(e)** Membrane binding of PEPA and PDPA (red) after incubation with A549 cells at 4 °C for 15 min. Scale bar = 20 μm. **(f, g)** The bio-nano interaction between the nanophotosensitizers and artificial lipid membrane measured by ITC at pH 6.0. **(h)** Interaction between endocytic organelle membrane (green) and Nile red-labelled PEPA and PDPA (red) visualized by HIS-SIM super-resolution microscopy. Scale bar = 10 μm. **(i)** Cell viability curves of PEPA and PDPA-mediated early endosome stress (EE stress at 0.5 h after internalization of nanoparticle) against A549 cells measured by MTT assay. **(j)** Immunoblots of PLC-G1 phosphorylation and GSDME cleavage of A549 cells treated with PEPA_EE_ and PDPA_EE_. PLC-G1, phospholipase C-γ1; p-PLC-G1, phosphorylated PLC-G1; GSDME-FL, full-length of GSDME; GSDME-N, N-terminal of GSDME. **(k)** Schematic of endo-lysosome stress controls the pyroptosis and apoptosis of cancer cells. Cells were pulsed with PEPA for 30 min and then chased at 37 °C for endosome maturation. 660 nm irradiation (6 J cm^−2^) was carried out at 0 h and 2 h for EE and lysosome (Ly) stress, respectively. **(l)** Representative phase-contrast images of CT26 cell death morphology induced by PEPA_EE_ or PEPA_Ly_ stress. **(m)** Cell viability curves of PEPA_EE_ and PEPA_Ly_ stress against CT26 cells measured by MTT assay. **(n)** Half-maximal inhibitory concentration (IC50) of PEPA_EE_ and PEPA_Ly_ stress against a series of murine cancer cell lines. **(o)** Immunoblots of GSDME cleavage of various murine cancer cells treated with PEPA_EE_ stress. Data are presented as mean ± s.d. (*n* = 3 biologically independent experiments).

We then chose PEPA as the top candidate of EE-targeted nanophotosensitizer to investigate the endosome maturation-gated pyroptosis on murine cancer cells (**Fig. 2k**). As indicated in **Extended Fig. 2a**, PEPA elicited precise oxidative stress in early endosomes (PEPA_EE_) and lysosomes (PEPA_Ly_) at 0.5 h and 2 h post-endocytosis, respectively. Effective cell killing with typical pyroptotic morphologies was evoked by PEPA_EE_ stress, while PEPA_Ly_ stress induced deficient apoptosis on CT26 colorectal cancer cells with a 4.9-fold higher IC_50_ value (**Fig. 2l,m** and **Supplementary Video 1**). Genetic knockout of

GSDME studies confirmed that PEPA achieved switchable pyroptosis/apoptosis on CT26 cells through endosome maturation (**Extended Fig. 2b,c**). We further generalized the PEPA-mediated tunable pyroptosis to a series of murine cancer cell lines with diverse expression level of GSDME (**Extended Fig. 2d, e**). Intriguingly, among the 8 cancer cell lines expressing GSDME, all demonstrated a range of 2.6- to 16.7-fold tunable photocytotoxicity between PEPA_EE_ and PEPA_Ly_, with a discrepancy of 3.8- to 5.0-fold in IC_50_ values (**Fig. 2n** and **Extended Fig. 2f**). Additionally, robust GSDME-mediated pyroptosis was observed on all cell lines upon treatment with PEPA_EE_ stress (**Fig. 2o** and **Extended Fig. 2g**).

### EE stress-induced pyroptosis amplifies tumour immunogenicity

We next investigated whether the organelle stress-mediated tunable pyroptosis can manipulate the tumour immunogenicity (**Fig. 3a**). Stressed and dying cancer cells release numerous bioactive molecules, including antigens and DAMPs. PEPA_EE_-stressed CT26 cells showed a 4.05-fold increase in protein release and a more diverse of types, as compared to the cells under PEPA_Ly_ stress. Notably, knockout of GSDME efficiently blocked the PEPA_EE_-mediated cellular content release (**Extended Fig. 3a,b**). Proteomics studies revealed that more released protein copies were induced by PEPA_EE_ (**Fig. 3b** and **Supplementary Fig. 4a**). Among the 351 copies of PEPA_EE_-specific proteins, the majority were identified in the nuclei and cytoplasm (**Supplementary Fig. 4b**). Gene Ontology (GO) enrichment analysis revealed their enrichment in categories related to stress response and immune response (**Fig. 3c**). Regarding the altered proteins, significance analysis indicated that 6 and 1136 proteins were decreased and increased in the supernatant of PEPA_EE_-stressed cells, respectively (**Fig. 3d**). Clustering and KGEE enrichment analysis further identified that, in comparison with PBS and PEPA_Ly_, PEPA_EE_ significantly boosted the release of immunogenicity-associated proteins (tumour antigen, DAMPs) and immune response-associated signalling pathways, including recruitment of immune cells, antigen presentation, and innate immunity (**Fig. 3e** and **Supplementary Fig. 4c**). The amplification of immunogenicity by PEPA_EE_ was also confirmed by the 34.8- (HMGB1) and 56.2-fold (ATP) enhanced release of ‘find me’ signals, as well as 4.6- (Annexin V), 8.2- (HSP70), and 21.0-fold (CRT) elevated membrane exposure of ‘eat-me’ signals, respectively. By contrast, both PEPA_Ly_ and GSDME deficiency resulted in a modest increase in the immunogenicity of CT26 cells (**Fig. 3f** and **Extended Fig. 3c-f**).

**Fig. 3.**
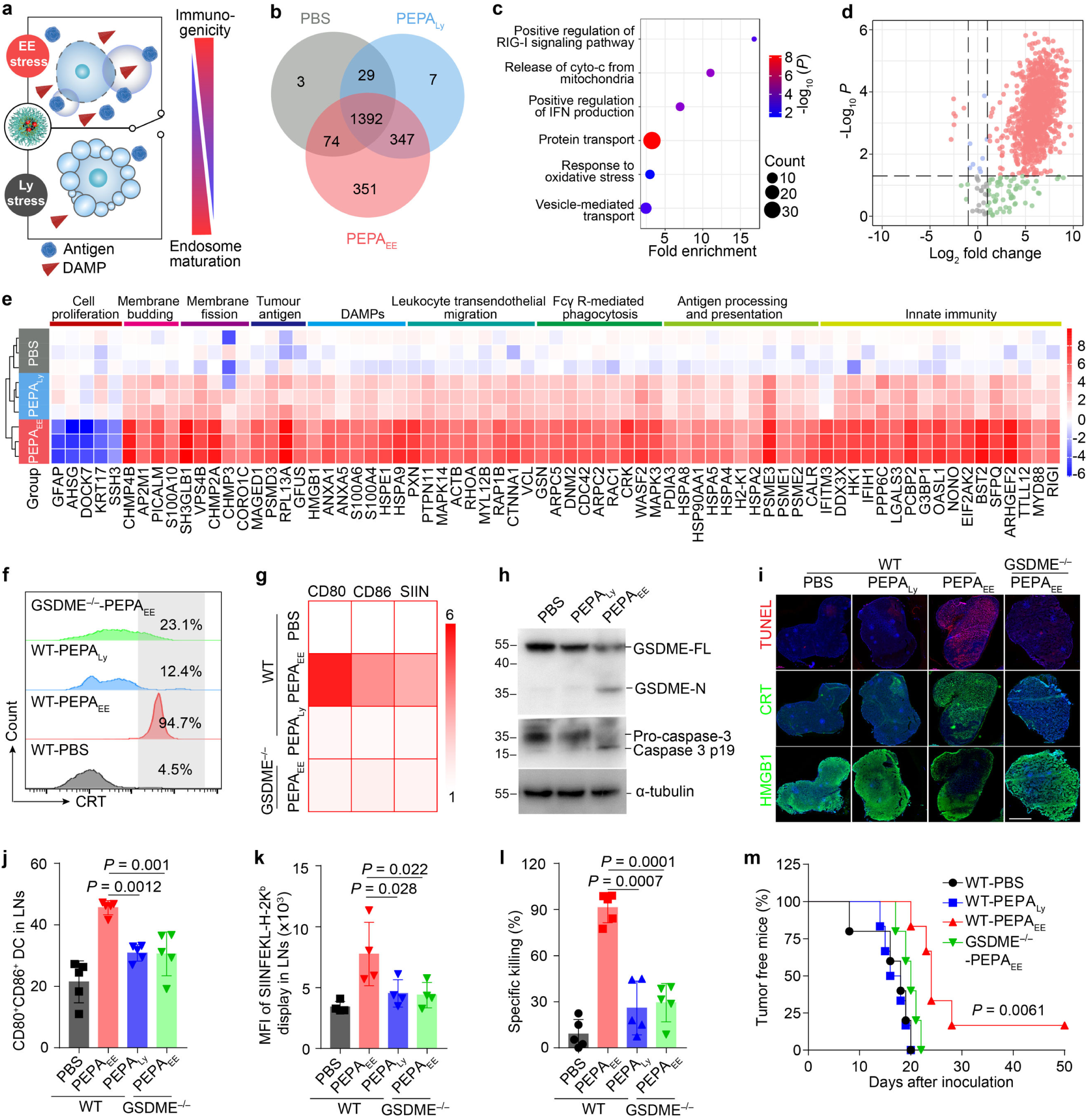
EE stress-evoked pyroptosis manipulates the immunogenicity of cancer cells. **(a)** Schematic illustration of PEPA-mediated endo-lysosome stress tunes the immunogenicity of cancer cells. DAMP, damage-associated molecular patterns. **(b-e)** Proteomics analysis of protein levels released from CT26 cells treated with PEPA-mediated EE or Ly stress. **(b)** Venn diagrams of released proteins. **(c)** Enrichment of biological process GOs specifically evoked by PEPA_EE_ stress. **(d)** Volcanic diagram of released proteins induced by PEPA-mediated EE and Ly stress. **(e)** Heatmap and clustering of significantly altered protein type between PEPA_EE_ and PEPA_Ly_. *n* = 3 biologically independent experiments. **(f)** Calreticulin (CRT) exposure of CT26 cells after treatment with PEPA_EE_ or PEPA_Ly_ stress. **(g)** Overexpression of costimulatory factors (CD80 and CD86) and presentation of OVA antigen (SIIN, SIINFEKL-H-2K^b^) on BMDCs after co-culture with PEPA_EE_ or PEPA_Ly_ treated CT26-OVA cells. Data were quantified by flow cytometry and normalized to PBS treatment. **(h)** Immunoblots of GSDME cleavage and caspase-3 activation in CT26-OVA tumours treated with PEPA_EE_ or PEPA_Ly_ stress. **(i)** Whole-slide imaging of immunogenicity (TUNEL, CRT exposure, and HMGB1 release) of tumour tissues elicited by PEPA_EE_ or PEPA_Ly_. Scale bar = 2 mm. **(j, k)** In vivo DC activation and presentation of OVA in draining lymph nodes harvested from CT26-OVA tumour-bearing mice after treatment with PEPA_EE_ or PEPA_Ly_. **(j)** Percentage of CD80^+^CD86^+^ DC cells, *n* = 5 mice. **(k)** Quantification of the SIINFEKL display among antigen-positive DCs, *n* = 4 mice. **(l)** Specific cell killing study in CT26-OVA tumour-bearing mice after different treatments (*n* = 5 mice). **(m)** Tumour occurrence in PEPA_EE_- or PEPA_Ly_-treated CT26 tumour-bearing mice after rechallenging with CT26 cells. *n* = 6 mice; log-rank test; *P* = 0.0061 for WT-PEPA_EE_ versus WT-PEPA_Ly_. All data are presented as mean ± s.d., and all measurements (*n*) are biologically independent.

To investigate the immune response of distinct stressed cancer cells, murine bone marrow-derived dendritic cells (BMDCs) were co-cultured with the dying CT26-OVA cells. The PEPA_Ly_-treated cells elicited negligible DC activation, showing minimal upregulation of co-stimulatory markers (CD80 and CD86), presentation of OVA (SIINFEKL), and secretion of cytokines (IL-12 and TNF-α). This is probably attributed to the immunologically silent nature of apoptosis^27^. Conversely, remarkable BMDC activation with a one-order-of-magnitude higher cytokine secretion was provoked by PEPA_EE_-stressed pyroptotic cells. As expected, PEPA_EE_-treated CT26-GSDME^−/−^ cells without pyroptotic phenotype exhibited deficient BMDC activation (**Fig. 3g** and **Extended Fig. 3g-j**). We further evaluated the organelle stress-mediated tunable immunogenicity and immune response in CT26-OVA tumour-bearing mice. Immune blotting and immunofluorescence staining studies demonstrated that PEPA_EE_ stress successfully evoked remarkable pyroptosis and ICD effect in tumour tissues in vivo (**Fig. 3h,i**). Consequently, a higher percentage of CD80^+^CD86^+^ DC cells with presentation of OVA antigen was observed in lymph nodes (LNs) of mice immunized with PEPA_EE_, in comparison with those in the PEPA_Ly_ treated or CT26-OVA-GSDME^−/−^ tumour-bearing mice (**Fig. 3j,k**). DC maturation can elicit the cytotoxic T cell-mediated adaptive immunity. In vivo specific cell killing studies indicated that 91.2% of OVA-positive splenocytes were specifically eliminated in PEPA_EE_-treated mice, while the percentages decreased to 26.0% and 29.4% in PEPA_Ly_-stressed mice and CT26-OVA-GSDME^−/−^ tumour-bearing mice, respectively (**Fig. 3l** and **Extended Fig. 3k**). The superior in vivo specific killing of PEPA_EE_ further enhanced the tumour prevention capacity when mice were rechallenged with CT26 cells, resulting in a 1.6-fold prolonged median relapse-free survival time (25 days) and suppressed tumour growth profiles (**Fig. 3m** and **Extended Fig. 3l,m**). Hence, we demonstrated that the immunogenicity of cancer cells can be modulated through elicitation of distinct endocytic organelle stress, and PEPA_EE_ stress-mediated pyroptosis efficiently boosted immunogenic cell death and cancer immune response in vitro and in vivo.

### Design of SPEN for safe and effective immune priming

A candidate vaccine requires both antigenicity and adjuvanticity^28^. While PEPA_EE_-elicited pyroptotic cells exhibit abundant antigenicity, the adjuvanticity of the released DAMP molecules is insufficient for efficient cancer vaccination. Hence, a TLR-7/8 agonist, IMDQ, was conjugated to the hydrophobic block of PEPA through thioketal linkage to construct the light-gated and ROS-sensitive SPEN nanoadjuvant (**Extended Fig. 4a**). PEPA conjugated with IMDQ through hydrocarbon linker was also synthesized to formulate the ROS-insensitive nanoadjuvant (SPIN). Synthesis and characterization of the polymer-drug conjugates were showed in **Supplementary Table 1** and **Supplementary Figs. 5-13**. The polymer-drug conjugates can self-assemble into nanoadjuvants. Upon 660 nm laser irradiation, more than 80% of total IMDQ released from SPEN, while no drug release from SPIN (**Extended Fig. 4b**). The light-gated immune activation capacity of SPEN were investigated on a TLR reporter cell line (Raw-Blue) and BMDC. As shown in **Extended Fig. 4c**, the 660 nm-irradiated SPEN exhibited robust TLR activating activity (EC_50_ = 85 nM) comparable to free IMDQ. Negligible TLR activation was observed on Raw-Blue cells treated with 660 nm-irradiated SPIN or SPEN without irradiation. Furthermore, 660 nm irradiation switched on the BMDC activation capacity of SPEN, exhibiting a binary pattern of membrane expression of CD80 and CD86, as well as secretion of IL-12 and TNF-α (**Extended Fig. 4d-g**). Importantly, SPEN efficiently abolished the undesired systemic and immune toxicity evoked by intravenous injection of free IMDQ, as verified by the minimized body weight loss, splenomegaly, and cytokine storm (**Extended Fig. 4h-m**). Therefore, the light-gated design of SPEN enables systemic delivery of immune adjuvant for safe and tumour-specific immune priming.

**Fig. 4.**
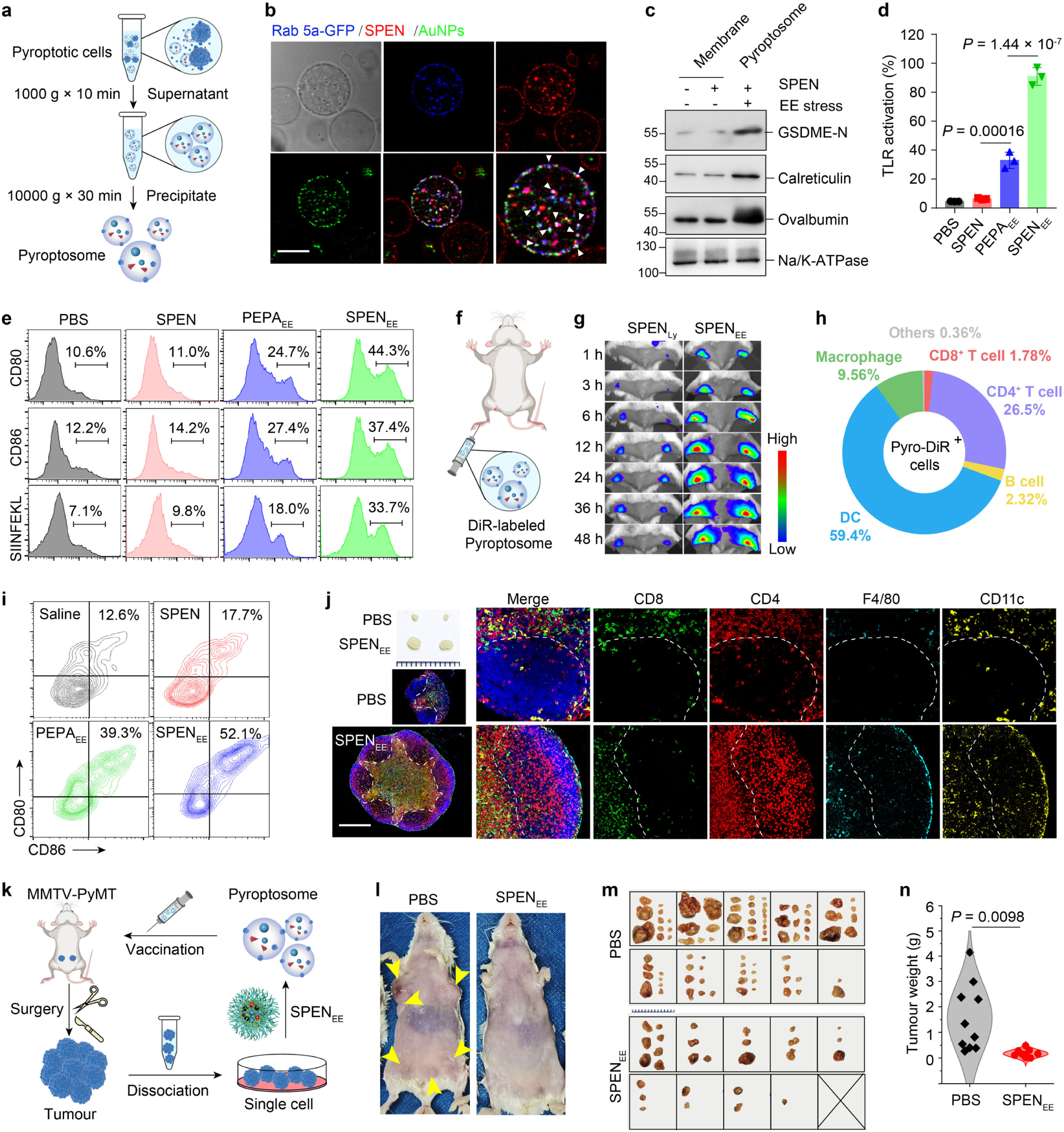
SPEN_EE_-derived pyroptosome functions as cancer vaccine. **(a)** Purification of pyroptosomes from pyroptotic cancer cells after treatment under SPEN_EE_ stress. **(b)** Confocal images of pyroptosomes from CT26 cells after treatment with SPEN_EE_ stress. CT26 cells were transfected with Rab5a-GFP, and a 10 nm Au nanoparticle (AuNP) was encapsulated in SPEN. Scale bar = 500 nm. **(c)** Immunoblots of GSDME-N, calreticulin, and OVA in pyroptosomes and cell membrane. **(d)** TLR activation efficacy of pyroptosomes from CT26 cells treated with PEPA_EE_ or SPEN_EE_ stress via Raw-Blue reporter cell assay. Data are shown as mean ± s.d. (*n* = 3 biologically independent experiments). **(e)** Overexpression of costimulatory factors (CD80 and CD86) and presentation of OVA antigen (SIINFEKL-H-2K^b^) on BMDCs after incubation with pyroptosomes from CT26-OVA cells. **(f)** BALB/c mice were immunized with pyroptosomes subcutaneously at foot pads for cancer vaccination. **(g)** In vivo fluorescence imaging of LN draining of pyroptosomes. Mice were injected with DiR-labelled pyroptosomes or SPEN_Ly-_treated CT26 cells on foot pads. **(h)** The proportions of various immune cells among the pyroptosome-DiR^+^ cells inside LNs. **(i)** Representative contour plots of CD80^+^CD86^+^ DC subsets among CD11c^+^ DCs in LNs of mice immunized with different pyroptosomes. **(j)** Representative photographs of LNs (scale bar = 2 mm) and corresponding sections after Opal multiplex staining with CD4 (CD4^+^ T cell, red), CD8 (CD8^+^ T cell, green), CD11c (DC, yellow), and F4/80 (macrophage, cyan) antibodies. The paracortical T cell regions are indicated by the white dashed lines. **(k-n)** Personalized cancer vaccination of SPEN-derived pyroptosomes on MMTV-PyMT model. **(k)** MMTV-PyMT/FVB mice without tumour occurrence were immunized with pyroptosomes prepared from spontaneous tumours of MMTV-PyMT/FVB mice. **(l)** Representative photographs of MMTV-PyMT mice immunized with PBS or SPEN_EE_-derived pyroptosomes. **(m)** Photographs of excised mammary tumour tissues from immunized MMTV-PyMT mice. Scale bar = 1 mm. *n* = 10 biologically independent mice. **(n)** The weights of excised mammary tumours. Data are shown as mean ± s.d.; *n* = 10 biologically independent mice; two-sided Student’s t-test with Welch’s correction.

### SPEN-mediated EE stress generates pyroptosome as an effective cancer vaccine

Cancer cell membranes have been widely applied as carriers and antigen source for cancer vaccine^11^. Then, we investigated the vaccine efficacy of pyroptotic cells provoked by SPEN-mediated EE stress (SPEN_EE_), focusing particularly on the bubbling pyroptotic membranes. After purification with differential centrifugation (**Fig. 4a**), we obtained the micro-sized pyroptosomes (**Extended Fig. 5a**). To investigate the composition of pyroptosomes, Rab5a-GFP transfected CT26-OVA cells were stressed with a 10 nm Au nanoparticle (AuNP)-encapsulated SPEN (**Extended Fig. 5b**). We observed an enrichment of the early endosomes, internalized SPEN, and IMDQ in the pyroptosomes (**Fig. 4b** and **Extended Fig. 5c**). Immune blotting of the membranes further revealed that the cleaved GSDME, CRT, and OVA in the pyroptosomes were 5.1-, 2.5-, and 2.9-fold higher than those in the untreated cell membranes, respectively (**Fig. 4c** and **Extended Fig. 5d**). Subsequently, we explored the in vitro and in vivo vaccination of pyroptosomes. Benefited from the superior TLR stimulation activity and higher adjuvanticity (**Fig. 4d**), SPEN_EE_ significantly amplified the BMDC maturation and antigen presentation capacity of EE stress-elicited pyroptosomes (**Fig. 4e** and **Extended Fig. 5e-i**). Upon injection into the foot pads, pyroptosomes exhibited rapid, high, and long-lasting lymph node draining capacity. A 6.3-fold higher fluorescent signals were observed in the popliteal LNs of mice injected with DiR-labelled pyroptosomes as compared to those from Ly-stressed cells (**Fig. 4f,g** and **Extended Fig. 5j-l**). Flow cytometry studies further demonstrated that 69.0% of DiR-positive cells were DCs and macrophages, indicating that the majority of pyroptosomes were internalized by antigen-presenting cells (APCs, **Fig. 4h**). Immunization with pyroptosomes further contributed to lymphadenectasis with 4.0-fold increase in weight. The percentages of matured DCs in LNs from mice immunized with SPEN_EE_- or PEPA_EE_-elicited pyroptosome were 4.1- and 3.1-fold higher than those from naïve mice, respectively, accompanied by a significantly elevated presentation of OVA (**Fig. 4i**, **Extended Fig. 5m,n**, and **Supplementary Fig. 14**). The immune response in LNs was further verified by the elevated proliferation of immune cells, infiltration of APCs in the paracortical T cell zone, and migration of CD4^+^ T cells into germinal center (**Fig. 4j** and **Extended Fig. 5o,p**). These results demonstrated that pyroptosomes can effectively deliver the tumour antigens and adjuvants to the LNs, functioning as a cancer vaccine with high immunogenicity.

**Fig. 5.**
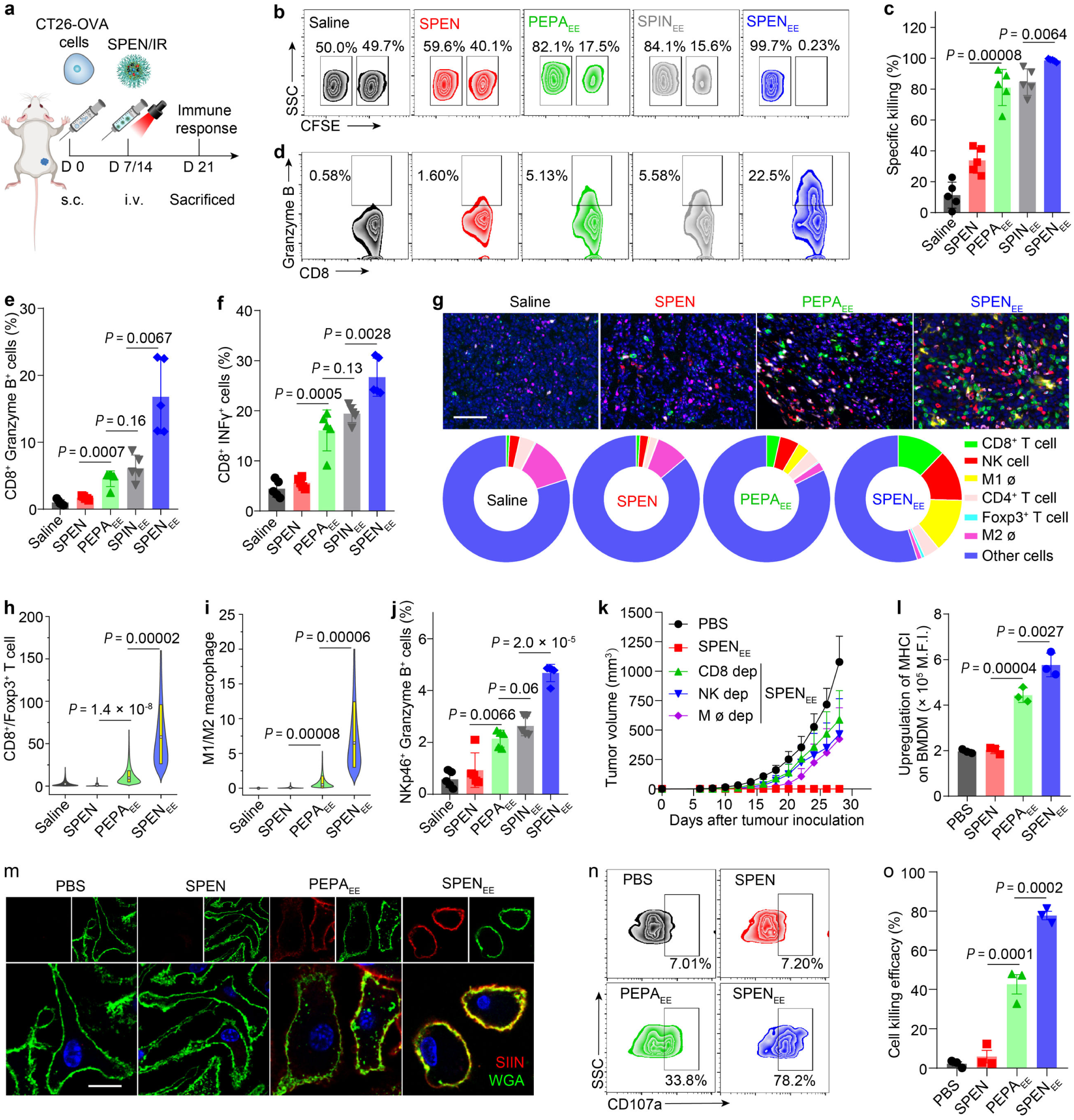
SPENEE in situ reprograms the anti-tumour immune microenvironment. **(a)** Schematic illustration for SPEN_EE_-mediated in situ vaccination on CT26-OVA tumour. CT26-OVA tumour-bearing mice were treated with SPEN-mediated EE stress every week for twice, and immune response studies were evaluated one week after the last treatment. **(b, c)** In vivo OVA-specific killing efficacy of various nanoadjuvants in CT26-OVA tumour-bearing mice, shown by representative flow cytometry plots **(b)** and quantification analysis as normalized to PBS treated mice **(c)**. *n* = 5 biologically independent mice. **(d, e)** The percentage of granzyme B^+^ cytotoxic T cells among CD8^+^ T cells in immunized mice. *n* = 5 biologically independent mice. **(f)** The percentage of interferon-γ (INF-γ)^+^ cytotoxic T cells among CD8^+^ T cells in immunized mice. *n* = 5 biologically independent mice. **(g)** Opal multiplex immunohistochemistry staining of tumour slides from immunized mice and corresponding pie chart showing the proportions of various lymphocytes in tumour microenvironment. Slides were immunostained with CD4 (CD4^+^ T, pink), CD8 (CD8^+^ T, green), NKp46 (NK, red), iNOS (M1 ø, yellow), CD206 (M2 ø, magenta), and Foxp3 (Treg, cyan). Scale bar = 100 μm. **(h, i)** The ratio of CD8^+^ T cells to Foxp3^+^ Treg cells **(h)** and M1-like macrophages to M2 phenotype **(i)** quantified from images in panel **g**. *n* = 20-30 biologically independent regions. **(j)** Percentage of intratumoural granzyme B^+^ NK cells among NKp46^+^ cells in immunized mice. *n* = 5 biologically independent mice. **(k)** Tumour inhibition of SPEN_EE_-mediated vaccination in CT26 tumour-bearing mice after depletion of CD8^+^ T cells, NK cells, or macrophages, respectively. *n* = 5 biologically independent mice. **(l)** Expression of MHC-I on cell membrane of BMDM after treatment with various pyroptosomes. *n* = 3 biologically independent experiment. **(m)** Confocal images of OVA presenting on BMDMs treated with various pyroptosomes. Scale bar = 20 μm. **(n)** NK activation by various pyroptosomes as indicated by the membrane expression of CD107a. **(o)** In vitro CT26 cell killing capacity of activated NK measured by LDH assay. *n* = 3 biologically independent experiment. All data are shown as mean ± s.d. Statistical significance was analyzed by one-way ANOVA followed by Dunnett’s multiple comparisons test.

Afterwards, we investigated the cancer prevention potency of pyroptosome-based vaccine. Immunization with SPEN_EE_-elicited pyroptosomes derived from CT26 cells successfully inhibited the growth of CT26 tumours in a dose-dependent manner. Tumour volume reduced to 1.6% when BALB/c mice were treated with pyroptosomes derived from 2 × 10^5^ pyroptotic cells, with a tumour incidence of 60% at day 22 post-inoculation of tumour cells (**Extended Fig. 5q** and **Supplementary Fig. 15**). However, the cancer vaccine efficacy of pyroptosomes was abolished in immunodeficient nude mice (**Supplementary Fig. 16**). Autologous tumours have been applied to produce cancer vaccines for personalized cancer immunotherapy to prevent post-operative recurrence and metastasis^7^. Hence, we excised the spontaneous breast tumours from MMTV-PyMT/FVB mice and applied the SPEN_EE_-based pyroptotic vaccine strategy (**Fig. 4k**). After immunization with autologous tumour-derived pyroptosomes, the development of mammary hyperplasia and neoplasia in transgenic mice was obviously suppressed (**Fig. 4l,m** and **Extended Fig. 5r**). Without pyroptosome vaccination, each mouse developed approximately 8.5 tumours with a total weight of 1.6 g. In pyroptosome-immunized mice, an average of 2.8 tumours (32.9%) with a relative low total weight (0.17 g, 10.6%) were observed in each mouse (**Fig. 4n** and **Extended Fig. 5s**).

### Pyroptosome spatiotemporally boosts the cancer-immunity cycle

Having confirmed the immunization functions of pyroptosomes, we next elucidated the underlying mechanism of SPEN-mediated vaccination (**Fig. 5a**). In vivo specific killing studies showed that the CT26-OVA tumour-bearing mice under treatment with PEPA_EE_ or SPIN-mediated EE stress (SPIN_EE_) exhibited OVA-specific cell killing of 81.1% and 84.6%, respectively. With the immune priming of IMDQ adjuvant, SPEN_EE_ stress achieved 99.7% OVA-specific cell killing efficacy (**Fig. 5b,c**). Compared with the negligible cytotoxic CD8^+^ T cells in naïve mice, the populations of granzyme B- and INF-γ-expressing cytotoxic CD8^+^ T cells were dramatically elevated in all EE stress-treated mice (**Fig. 5d** and **Extended Fig. 6a**). The percentages of granzyme B^+^/CD8^+^ T cells and INF-γ^+^/CD8^+^ T cells in mice immunized with SPEN_EE_ stress were boosted to 16.7- and 6.0-fold higher than those in naïve mice, respectively (**Fig. 5e,f**). These results revealed that SPEN_EE_ stress-mediated pyroptosis initiated the adaptive T cell immune response in vivo. To elucidate the intratumorally immunologic mechanism, we further visualized the tumour immune microenvironment (TIME) by multicolor fluorescence immunohistochemistry. As shown in **Fig. 5g**, SPEN_EE_ stress successfully reprogrammed the TIME, the tumour-suppressive immune cell populations, CD8^+^ T cells, natural killer (NK) cells, and M1-type macrophages increased to 20.3-, 14.8-, and 35.2-fold, respectively (**Extended Fig. 6b-d**). Conversely, the percentage of tumour-supportive M2-type macrophages decreased by 90% (**Extended Fig. 6e**). The significant expansion of CD8^+^-to-Foxp3^+^ T cell and M1-to-M2 macrophage ratios further elucidated that SPEN_EE_-mediated in vivo vaccination can reverse immune suppression within the tumour microenvironment (**Fig. 5h,i**). The cytotoxic functions of intratumoral NK cells were also verified by the amplified secretion of granzyme B and INF-γ on the effector NK cells (**Fig. 5j** and **Extended Fig. 6f,g**). Macrophages and NK cells play critical roles in innate immunity. To investigate the contribution of innate and adaptive immunity in SPEN_EE_-based immunization, we depleted different immune cells in CT26 tumour-bearing mice^10^ (**Supplementary Fig. 17**). Immunization with SPEN_EE_-mediated in situ vaccine achieved 100% cancer prevention, while depletion of CD8^+^ T cells, NK cells, and macrophages resulted in fast tumour relapse in immunized mice, respectively (**Fig. 5k**).

**Fig. 6.**
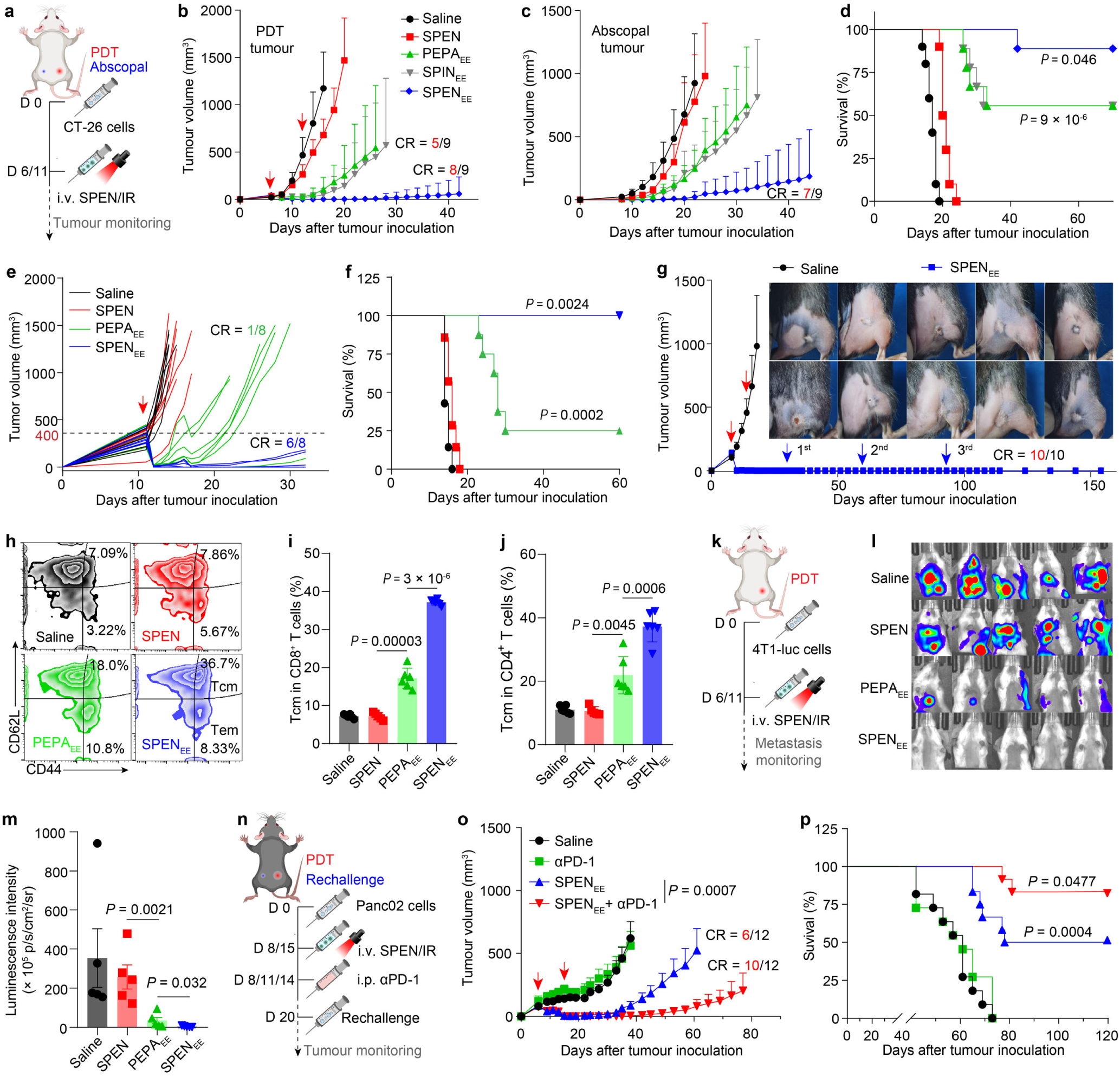
SPENEE achieves robust tumour immunotherapy and long-lasting vaccination against multiple cancers. (a-d) Tumour immunotherapy of SPEN-mediated EE stress on CT26 tumour-bearing mice (*n* = 10 mice for saline and SPEN groups; *n* = 9 mice for PEPA_EE_, SPIN_EE_, and SPEN_EE_ groups). Schematic illustration of experiment schedule **(a)**. Growth profiles of primary tumour with the treatment of EE stress **(b)**. Growth profiles of abscopal tumour without EE stress **(c)**. CR, completely removal. Animal survival curves **(d**; log-rank test; *P* = 9 × 10^-6^ for PEPA_EE_ versus SPEN, *P* = 0.046 for PEPA_EE_ versus SPEN_EE_). **(e, f)** Therapeutic efficiency of SPEN_EE_ on B16 tumour-bearing mice with large tumours of ∼400 mm^3^ (*n* = 7 mice for saline and SPEN groups; *n* = 8 mice for PEPA_EE_ and SPEN_EE_ groups). Individual tumour growth curves **(e)**. Animal survival curves **(f**; log-rank test; *P* = 0.0002 for PEPA_EE_ versus SPEN, *P* = 0.0024 for PEPA_EE_ versus SPEN_EE_). **(g-j)** Long-lasting cancer prevention capacity of SPEN_EE_ on MC38 tumour-bearing mice. Photographs of completely cured MC38 tumours and tumour growth profiles of SPEN_EE_-immunized mice rechallenged with MC38 cells for three time (**g**; *n* = 10 biologically independent mice). Representative scatterplots of CD44^high^ CD62L^low^ T cell (effector memory T cells, Tem) and CD44^high^ CD62L^high^ T cell (central memory T cells, Tcm) subsets among CD8^+^ T lymphocytes **(h)**. Quantification of Tcm in CD8^+^ T cells **(i)** and CD4^+^ T cells **(j)**. *n* = 6 biologically independent mice; one-way ANOVA followed by Dunnett’s multiple comparisons test. **(k-m)** Lung metastasis inhibition of SPEN_EE_ on 4T1-luc tumour-bearing mice. Schematic illustration of experiment schedule **(k)**. In vivo lung metastasis visualized by luminescence imaging **(l)**. Quantification of luminescence imaging of excised lungs **(m)**. One-way ANOVA followed by Dunnett’s multiple comparisons test. **(n-p)** Combination immunotherapy of SPEN with αPD-1 on the immunosuppressive PANC02 tumour-bearing mice (*n* = 11 mice for saline and αPD-1 groups; *n* = 12 mice for SPEN_EE_ and SPEN_EE_ + αPD-1 groups). Schematic illustration of experiment schedule **(n)**. Growth profiles of PANC02 tumour with various treatments (**o**; two-way ANOVA followed by Bonferroni’s multiple comparisons test). Animal survival curves **(p**; log-rank test; *P* = 0.0004 for αPD-1 versus SPEN_EE_, *P* = 0.0477 for SPEN_EE_ versus SPEN_EE_ + αPD-1). All data are shown as mean ± s.d.

We next investigated the functions of pyroptosomes on macrophages and NK cells in vitro. Macrophages represent one of the major APCs. Previous studies have verified that IMDQ-based nanoadjuvants selectively reprogramed the M2-type antigen-destroying tumour-associated macrophages to M1 phenotype with improved antigen presentation^24^. We also confirmed that SPEN_EE_-mediated pyroptosomes and released IMDQ profoundly improved the membrane presentation of MHC-I molecules and OVA antigens on M2-type macrophages (**Fig. 5l,m** and **Extended Fig. 6h,i**). Furthermore, the spleen-derived immature NK cells were successfully activated by pyroptosomes into effector NK cells with cytotoxic functions, as evidenced by the heightened membrane expression of NKp46 and CD107a (**Fig. 5n**, **Extended Fig. 6j,k** and **Supplementary Fig. 18**), resulting in 77.7% NK cell-mediated cytotoxicity towards CT26 cells (**Fig. 5o**). Collectively, these results demonstrated that SPEN-mediated in situ vaccination activated both innate and adaptive cancer immunity to close the cancer-immunity cycle.

### SPEN_EE_-mediated in situ vaccination achieves robust cancer immunotherapy

Given the satisfactory immunization capacity, we exploited SPEN_EE_ for in vivo cancer immunotherapy. First, the abscopal effect was assessed on CT26 cancer model (**Fig. 6a**). Results revealed that SPEN_EE_ stress remarkably eradicated the irradiated tumours and the abscopal ones with tumour-free rate of 88.9% (8/9) and 77.8% (7/9), respectively. Without the adjuvanticity of IMDQ, the efficacy of PEPA_EE_ and SPIN_EE_ was compromised, especially for the abscopal effect with tumour-free rate of 11.1% (1/9) and 0% (0/9), respectively (**Fig. 6b,c** and **Extended Fig. 7a**). The survival time of tumour-bearing mice treated with SPEN_EE_ was also significantly prolonged (**Fig. 6d**). Next, we evaluated the therapeutic efficacy on B16 melanoma model with large tumours of ∼400 mm^3^. While the majority of tumour burden was eradicated by PEPA_EE_ stress, 87.5% of mice experienced rapid tumour relapse during the withdrawal period. In comparison, 75% of large tumour-bearing mice were completely cured by SPEN_EE_, with no instances of animal death and LN metastasis during the two-month treatment-free period (**Fig. 6e,f** and **Extended Fig. 7b,c**).

To further certify the prophylactic and therapeutic activity of SPEN_EE_-mediated in situ vaccine, MC38 colorectal tumour-bearing mice were immunized with SPEN_EE_ stress (**Extended Fig. 7d**). The immunized mice achieved prominent anti-tumour efficacy, with 100% (10/10) of tumour regression. Importantly, all the tumour-free mice in SPEN_EE_ group successfully prevented multiple MC38 tumour rechallenge (day 30, day 60, and day 90) during the five-month treatment-free period (**Fig. 6g** and **Extended Fig. 7e,f**). The long-term cancer prevention beyond 150 days demonstrated that SPEN_EE_-mediated in situ vaccine evoked robust immunological memory. This was then corroborated by a 5.1- and 3.4-fold increase in central memory CD8^+^ T cells and CD4^+^ T cells, respectively (**Fig. 6h-j** and **Extended Fig. 7g**), as well as a significant elevation in effector memory T cells (**Supplementary Fig. 19**), compared to the saline group.

Benefited from its superior vaccination efficacy, SPEN_EE_ stress also noticeably circumvented lung metastasis in the highly invasive 4T1 tumour model, as indicated by the undetectable luminescence signals (eq. to 1.0% of saline group) in lung tissues (**Fig. 6k-m** and **Extended Fig. 7h-k**). Considering the poor efficacy of immune checkpoint blockade therapy on the cold tumours with immune-silent phenotype, we combined the SPEN_EE_-mediated in situ vaccine strategy with αPD-1 to potentiate therapeutic efficacy on Panc02 pancreatic cancer model (**Fig. 6n**). Negligible tumour inhibition was observed in mice with αPD-1 monotherapy, whereas 50% of the SPEN_EE_-immunized mice exhibited no tumour progression. Notably, combination therapy of SPEN_EE_ and αPD-1 achieved 83.3% eradiation of primary tumour, accompanied by significantly prolonged mice survival and cancer prevention within 120 days (**Fig. 6o,p** and **Extended Fig. 7l-n**). Altogether, the EE stress-gated immunogenic pyroptosis and adjuvant platform offers a promising in situ vaccine strategy for precise cancer immunotherapy.

## Discussion

Weak immunogenicity and poor availability of patient-specific tumour antigens are major obstacles that hamper the clinical therapeutic outcomes of cancer vaccine^3, 29^. In situ cancer vaccines, utilizing autologous tumour antigens at tumour sites, have been proposed as personalized cancer immunotherapy post-surgery^11^. To optimize the immunogenicity of in situ tumour vaccines, various strategies, including chemotherapy^30^, radiotherapy^31^, phototherapy^32^, and oncolytic virotherapy^33^ have been employed as cellular stress inducers for triggering the immunogenic cell death. However, the immunological correlates between cancer immunity and cellular stress-induced ICD effect, as well as the underlying immunogenic mechanism remains unclear. Benefiting from our spatiotemporally tunable pyroptosis nanotechnology, we succeeded to uncover the organelle stress-dependent amplification of cancer cell immunogenicity in vitro and in vivo.

Adjuvants play crucial roles in modulating the cold and suppressive tumour immune microenvironment to boost vaccine-induced immune response. TLR7/8 agonist, IMDQ, has been exploited as one of the most efficient chemical adjuvants in vaccine^34, 35^. However, the always-on immunostimulatory bioactivity and rapid diffusion of free IMDQ have resulted in serious adverse events, such as headache and cytokine storm, restricting its clinical application via subcutaneous and intravenous administration^36^. Increasing evidence has revealed that systemic delivery of agonists can intensify the exposure of dying tumour cells to adjuvants, thereby functioning as an in situ vaccine with superior therapeutic outcomes^37^. Therefore, we developed the pH/oxidative stress logical AND-gate nanoadjuvant, where IMDQ is chemically trapped in nanoparticles with quenched immunostimulatory bioactivity in non-malignant tissues, while achieving precise release in active form along with the formation of pyroptosome from EE-stressed cancer cells.

Our studies revealed that the SPEN nanoadjuvants with high membrane affinity can precisely elicit robust pyroptotic cancer cell death in early endosomes, and further generate pyroptosomes in situ with vigorous immunogenicity, whereas the lysosome stress provokes low immunogenic apoptosis. The photo-induced pyroptosomes serve as the source of plenty tumour antigens for polyclonal immune response; and the pyroptotic cell death triggers the release of multiple DAMPs (e.g. HMGB1, ATP, and CRT) and pro-inflammatory cytokines with up to 50-fold increment, thereby efficiently modulating the antigen presentation and immunosuppressive tumour microenvironment for the efficient cascade activation of CIC. Thus, the SPEN design enables the formation of tumour antigen-rich pyroptosomes and the release of active adjuvant in the right time (following pyroptosis of cancer cells) and at the right place (triggers the release of tumour antigen and adjuvant from the same tumour cells), serving as in situ cancer vaccines to boost the immune microenvironment of LNs and tumours, facilitate effective tumour immunity and long-term cancer prevention.

In summary, we have developed a systemic injectable and pyroptosis-enabled nanoadjuvant that works as an in situ cancer vaccine through boosting the cancer-immunity cycle. This study evidences the promising immunogenic functions of early endosome-stressed pyroptotic cell death, revealing the underlying immune-priming mechanisms of pyroptosis-derived vesicles, but also opens new opportunities to engineer pyroptosis-inducing nanomaterials for personalized cancer immunotherapy.

## Methods

### Cell lines and culture

A549 lung carcinoma (1101HUM-PUMC000002), 4T1 breast cancer (3101MOUSCSP5056), CT26 colon cancer (1101MOU-PUMC000275), and PANC02 pancreatic cancer (CRL-2553) cell lines were purchased from National Infrastructure of Cell Line Resource (Beijing, China). RAW-Blue and RAW-Blue ISG reporter cells were obtained from InvivoGen. MC38 colorectal cancer cells and luciferase-transfected 4T1 (4T1-Luc) cells were gifts from Dr. Wei Liang (Institute of Biophysics, Chinese Academy of Sciences). B16-F10 melanoma cells were kindly provided by Dr. Jinming Gao (University of Texas Southwestern Medical Center). ID8 ovarian cancer cell line was a gift from Dr. Yi Li (Peking University People’s Hospital). The CT26-GSDME^−/−^, CT26-GFP, and CT26-OVA cells were constructed by CRISPR-Cas9 technology as described previously^18^. All cells were cultured in DMEM medium with 10% fetal bovine serum (FBS) and antibiotics (penicillin 100 Uml^−1^ and streptomycin 100 μg mL^−1^) at 37 °C with 5% CO_2_. All cells were tested to be negative for mycoplasma.

### Animals and tumour models

Female BALB/c, C57BL/6, and FVB mice (6-8 weeks) were obtained from Vital River Laboratory Animal Center (Beijing, China). The MMTV-PyMT/FVB transgenic mouse breast cancer model was sourced from the Jackson Laboratory. The mice were housed at a temperature of 25 °C in a humidity-controlled environment with free access to food and water, and maintained in specific pathogen-free (SPF) conditions for one week before the studies. All care and handling of animals were approved by the Institutional Animal Care and Use Committee (IACUC) of Peking University (Accreditation number: LA 2023056). To establish tumour-bearing mice models, CT26 (1 × 10^6^), 4T1 (1 × 10^6^), B16 (5 × 10^5^), MC38 (2 × 10^6^), or PANC02 (2 × 10^6^) cells were subcutaneously injected on the right flank of BALB/c or C57BL/6 mice, respectively. The greatest longitudinal diameter (length) and transverse diameter (width) of tumours were measured by a vernier caliper. Tumour volume was calculated by the formula: (length×width^2^)/2. The maximum tumour volume did not exceeded 1,500 mm^3^ in any of the studies.

### Preparation and characterization of organelle-stressed nanophotosensitizer

The endocytic organelle-stressed nanoparticles were fabricated by the solvent displacement method as reported previously^18^. Briefly, 2.0 mg of PPa-conjugated copolymers (copolymerized with or without IMDQ monomers) was dissolved in 0.2 mL of THF, and added dropwise into 4.0 mL of Milli-Q deionized water under sonication for 40 seconds. Then the THF was removed via micro-ultrafiltration tubes (100 kDa, Merck Millipore), and the suspension was concentrated to 5.0 mg mL^-1^ for further studies. For characterization, dynamic light scattering (Zetasizer Nano ZSP, Malvern Instruments) was applied to measure the hydrodynamic diameter and zeta potentials of nanoparticles at room temperature. Transmission electron microscope (JEM-1400, JEOL) was used to evaluate the morphology of nanoparticles in different pH conditions. An IVIS Spectrum imaging system (Lumina Series III, Perkin Elmer, Living Image 4.3.1) was involved to assess the acid-amplified fluorescence property of nanoparticles in 100 mM PBS buffers with different pH values using an autoexposure model with bandpass filters (λ_ex_/λ_em_ refers to the wavelengths of excitation and emission, 620 ± 10 nm/670 ± 20 nm).

### Subcellular trafficking

The subcellular endo-lysosomal colocalization studies of PEPA and PDPA were investigated by CLSM. The CellLight Early Endosome (Rab5a)-GFP and Lysosome (Lamp1)-GFP BacMam 2.0 kits (Molecular Probes) were applied to label the early endosomes (EE) and lysosomes (Ly) of A549 cells, respectively. Nuclei were labelled with Hoechst 33342 at 37 °C for 20 min. Then the cells were incubated with a mixture of PEPA-Cy5 and PDPA-TMR at 37 °C for 15 min. After being replaced with fresh medium, subcellular localization of nanoparticles was captured at predesignated time-points by confocal fluorescence microscopy. The DAPI, FITC, TRITC, and Cy5 filters were used for Hoechst 33342, GFP, TMR, and Cy5 imaging, respectively.

### Visualization of the affinity of protonated polymers with endo-lysosomal membrane

The Lysosome-GFP-labelled A549 cells were incubated with Nile red (NR)-conjugated PEPA or PDPA at a NR concentration of 1 μg mL^−1^ (diluted with culture medium) at 37 °C for 2 h, followed by nuclei labelling. After being replaced with fresh medium, the affinity between nanophotosensitizers and endo-lysosomal membranes was visualized with a HIS-SIM super-resolution microscope (CSR Biotech.).

### In vitro cell killing efficacy

To compare the photocytotoxicity of PEPA and PDPA nanophotosensitizers, A549 cells were seeded on a 24-well plate at a density of 2×10^4^ cells per well and incubated at 37 °C overnight before the assay. The culture medium was then replaced with fresh medium containing a series of PPa concentrations of the PEPA or PDPA nanophotosensitizers. After incubation at 37 °C for 0.5 h, the treated cells were irradiated with a 660 nm-laser (100 mWcm^−2^) for 1 min. The MTT assay was carried out to measure the cell viability at 24 h after irradiation. Briefly, 0.5 mL MTT stock solution was added to the wells at a final concentration of 0.5 mg mL^−1^, and the generated formazan was dissolved with DMSO solvent after 1 h incubation. The absorbance at 540 nm was quantified on a microplate reader (Multiskan FC, Thermo Fisher Scientific). The IC50 values of the nanophotosensitizers on A549 cells were calculated by Origin Pro software. The GFP-transfected A549 cells and PI (4.5 μM, Sigma) staining were included to evaluate the cell killing efficacy by CLSM.

To estimate the cell killing efficacy of EE and Ly stress mediated by PEPA nanophotosensitizer, a series of murine cancer cell lines, including CT26, 4T1, B16, MC38, TC-1, PANC02, ID8, and KPC, were treated with a series of PPa concentrations of the PEPA. Irradiation (100mWcm^−2^ for 1 min) was carried out at 0 h (PEPA_EE_) or 2 h (PEPA_Ly_) post-incubation with nanophotosensitizer, respectively.

### Cell death morphology imaging

To observe the cell morphology, cells were seeded in glass-bottom dishes at about 50% confluence and subjected to the indicated treatments for static image capture and living imaging. Representative bright-field images of cell death were observed and captured with a Zeiss Axio Vert AI or Zeiss LSM880 (ZEN 2010) confocal microscope. The images were processed with ImageJ 1.47 software. Live images of cell death after the indicated treatments were also recorded in real time by a Zeiss LSM880 confocal microscope and processed with Corel VideoStudio X10 program. Cells were transfected with GFP (CRISPR-Cas9 technology) to indicate the integrity of the cell membrane. All image data are shown as representative of at least three randomly selected fields.

### Immunoblot analysis

The cellular expression of α-tubulin, phosphorylation of PLC, and cleavages of GSDME were all detected by western blot analysis. Cells with different treatments were lysed in buffer containing 25 mM tris(hydroxymethyl) aminomethane (Tris; pH 7.4), 150 mM NaCl, 0.5% sodium deoxycholate, 0.1% SDS buffer, and 1% Triton X-100 surfactant, as well as cocktail protease inhibitor (Roche) and phosphatase inhibitor (Roche). Cell lysates were clarified by spinning at 13,800 g for 10 min at 4 °C, and supernatants were diluted with × 4 loading buffer, followed by boiling for 10 min. Denatured proteins (20 μg) were fractionated by sodium dodecyl sulfate–polyacrylamide gel electrophoresis and transferred to polyvinyl difluoride membranes with a Mini-PROTEAN Tetra system (Bio-Rad). Blots were then blocked with 5% skim milk in Tris-buffered saline plus 0.05% Tween-20 and probed with appropriate antibodies according to the manufacturer’s instructions.

### Shotgun proteomics

The CT26 cells treated with PBS, PEPA_EE_ or PEPA_Ly_ stress were centrifuged at 1,000 × g for 10 min. The supernatants were collected and lysed in 0.5% sodium deoxycholate containing with 25 mM Tris, 150 mM NaCl, 0.1% SDS, 1% Triton X-100, cocktail protease inhibitor and phosphatase inhibitor (Roche). Lysates were clarified by centrifugation at 13,800 × g for 10 min and quantified to be 200 μg protein from each sample, followed by trypsin (Promega) digestion overnight for subsequent liquid chromatography-mass spectrometry (LC-MS) analysis (*n* = 3).

### In vitro and in vivo immunogenic cell death

CT26 and CT26-GSDME^−/−^ cells were seeded in 24-well plates overnight, and treated with PEPA_EE_ or PEPA_Ly_. The supernatants were collected for SDS-PAGE and ATP release assay by ATPlite Luminescence Assay System (PerkinElmer). The treated cells were washed with PBS, trypsinized, and collected, followed by staining with an AF488-labelled anti-CRT antibody or AF488-labelled anti-HSP70 antibody for 30 min at 4 ℃. Then the membrane exposure of CRT and HSP70 were quantified by flow cytometry. To visualize the cell immunogenicity, CT26 and CT26-GSDME^−/−^ cells were seeded in glass-bottom dishes and subjected to the indicated treatments. Then the cells were stained with an Annexin V-FITC/PI Apoptosis Detection Kit (Beyotime) according to the manufacturer’s instructions. In HMGB1 release study, the treated cells were fixed with 4% paraformaldehyde, permeabilized by 0.1% Triton X-100, and blocked with 5% BSA in PBS for 1 h (r.t.). The cells were then incubated with Rabbit anti-HMGB1 antibody (1:500) overnight at 4 °C followed by Alexa Flour 594-conjugated Goat anti-Rabbit IgG staining for 2 h (r.t.). The stained cells were washed, labelled with Hoechst 33342, and imaged with CLSM (ZEISS LSM880).

For in vivo ICD evaluation, the CT26 and CT26-GSDME^−/−^ tumour-bearing mice were treated with PEPA_EE_ or PEPA_Ly_. The tumour tissues were excised, sectioned for in situ apoptosis analysis by TUNEL assay according to the manufacturer’s instructions. The adjacent tumour slides were fixed, and blocked for immune staining with anti-CRT and anti-HMGB1 antibodies, respectively. Then, the slides were sealed with Hoechst 33342-containing antifade mounting medium, and the whole-slide images were captured by a Vectra Polaris scanner (PerkinElmer).

### Preparation and characterization of pyroptosome

To purify the pyroptosomes, the pyroptotic cancer cells were centrifuged at 1,000 × g for 10 min to remove the living cells and cell fragments. The supernatant was then centrifuged at 10,000 × g for another 30 min. The pyroptosomes were obtained by resuspending the precipitate in PBS. The morphology of pyroptosomes was confirmed by TEM and CLSM. To investigate the composition of pyroptosomes, the Rab5a-GFP transfected CT26-OVA cells were stressed by a 10 nm Au nanoparticle (AuNP)-encapsulated SPEN, and the obtained pyroptosomes were visualized by CLSM. The pyroptosomes were further lysed and immunoblotted with anti-GSDME, anti-CRT, anti-Rab5a, anti-ovalbumin, and anti-Na/K-ATPase antibodies.

### In vitro activation of BMDC

Murine bone marrow-derived dendritic cells (BMDCs) were extracted and cultured as previously reported^38^. To investigate the immunogenicity of EE-stressed tumour cells, CT26-OVA and CT26-OVA-GSDME^−/−^ cells were seeded in 24-well plates, and treated with PEPA_EE_ or PEPA_Ly_. Then 2×10^5^ BMDCs were added into the treated tumour cells. To assess the light-gated DC activation of SPEN, BMDCs were seeded in 24-well plates, followed by treatments of different IMDQ formulations (1 μM IMDQ) with or without 660 nm irradiation. In another experiment, the BMDCs were treated with different EE-stressed and CT26-OVA-derived pyroptosomes. Following 24 h of incubation, BMDCs were collected and stained with FITC anti-mouse CD11c, PE/Cy7 anti-mouse CD80, APC anti-mouse CD86, and PE anti-mouse SIINFEKL-H-2K^b^ (Biolegend) to estimate the expression of co-stimulaters and presentation of OVA by flow cytometry. The supernatants were collected to quantify the secretion of IP-10 and IL-12 by ELISA kits (Peprotech).

### Lymph node draining and distribution of pyroptosomes

CT26 cells were incubated with DiR (5 μM) for 15 min at 37 °C to label the cell membrane, and treated with SPEN-mediated EE stress. Then the DiR-labelled CT26 cells were injected in the foot pad of BALB/c mice. In vivo fluorescence images of DiR signals were captured at predesignated time-points with IVIS Spectrum imaging system, and the fluorescence intensity of popliteal lymph nodes was quantified by Living Image software. At 48 h post-injection, the popliteal lymph nodes were excised for imaging. Afterwards, the LNs were prepared into single cell suspensions and immunostained with PerCP/Cy5.5 anti-mouse CD45, FITC anti-mouse CD3, PE anti-mouse CD4, APC anti-mouse CD8, BV421 anti-mouse CD11c (DC), PE anti-mouse CD19 (B cell), and APC anti-mouse F4/80 (macrophage) for flow cytometry analysis.

### Lymph nodes activation in vivo

CT26-OVA tumour-bearing BALB/c mice were intravenously injected with PEPA, SPEN, or SPIN at an equal PPa dose of 1.0 mg kg^−1^ and IMDQ dose of 0.4 mg kg^−1^. Mice were anaesthetized with 1.2% avertin, and the tumour sites were irradiated with a 660 nm laser (240 J cm^−2^) at 3 h post-injection (EE stress). The treatments were repeated once a week for twice. At day 14 after the first treatment, the inguinal and popliteal lymph nodes were excised, prepared into single cell suspensions for DC and T cell staining. For DC analysis, the cell suspensions were stained with FITC anti-mouse CD11c, PE/Cy7 anti-mouse CD80, APC anti-mouse CD86, and PE anti-mouse SIINFEKL-H-2K^b^. For T cell analysis, the cell suspensions were labelled with PerCP/Cy5.5 anti-mouse CD45, FITC anti-mouse CD3, PE anti-mouse CD4, and APC anti-mouse CD8. Then the cell suspensions were analyzed by flow cytometry.

### In vivo cancer prevention of pyroptososmes

BALB/c mice were subcutaneously vaccinated with SPEN_EE_-derived pyroptosomes from 0, 2×10^4^, 4×10^4^, 1×10^5^, or 2×10^5^ pyroptotic CT26 cells once a week for three times, respectively. The immunized mice were inoculated with 5×10^5^ CT26 cells (5×10^5^ cells/mouse) in the right flank seven days after the last vaccination. The tumour growth profiles were monitored to evaluate the cancer prophylactic efficacy of pyroptosomes. For MMTV-PyMT/FVB transgenic breast cancer model, tumour tissues were dissected from mice with spontaneous tumours, and prepared into single cell suspensions. Cells were treated with SPEN-mediated EE stress, and pyroptosomes were purified for vaccination. MMTV-PyMT/FVB mice (4-6 week) without tumour occurrence were immunized with personalized pyroptosomes (2×10^5^ pyroptotic cells per mice) weekly for three times. The vaccination efficacy was investigated by monitoring the mammary hyperplasia and carcinoma.

### In vivo specific cell killing assay

CT26-OVA tumour-bearing mice were intravenously injected with PEPA, SPEN, or SPIN at an equal PPa dose of 1.0 mg kg^−1^ and IMDQ dose of 0.4 mg kg^−1^. Irradiation (240 J cm^−2^) was carried out at 3 h post-injection (EE stress). The treatments were repeated once a week for twice. At day 14 after the first treatment, spleens from naïve mice were processed to single-cell suspension. The splenocytes were pulsed with OVA_257-264_ (10 μg/mL) or blank DMEM medium for 30 min, and labelled with 5 μM or 0.5 μM carboxyfluorescein succinimidyl ester (CFSE, BD) for 15 min, respectively. Then the OVA^+^ CFSE^high^ and OVA^-^ CFSE^low^ splenocytes were equally mixed (1×10^7^) and intravenously injected into the treated mice. After two days, the immunized mice were euthanized and splenocytes were collected for flow cytometry to evaluate the intracellular CFSE signal. Specific cell killing efficacy is calculated according to the following formula:

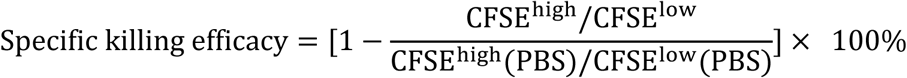

### Cytotoxic T cell and NK activation

CT26-OVA tumour-bearing mice were immunized with different nanophotosensitizers and EE stress. The tumours were dissociated into single cells, and cultured with cell stimulation cocktail (eBioscience) for 6 h at 37 °C. Then the cells were stained with APC anti-mouse CD8α or APC anti-mouse NKp46 antibody, respectively, followed by treatment with Intracellular Fixation & Permeabilization Buffer Set (eBioscience) according to the manufacturer’s instructions, and labelled with PE anti-mouse IFN-γ or PE anti-mouse Granzyme B (GranzB) antibody. The percentages of CD8α^+^IFN-γ^+^, CD8α^+^GranzB^+^, NKp46^+^IFN-γ^+^, and NKp46^+^GranzB^+^ cytotoxic cells were quantified by flow cytometry.

### In vitro activation and antigen presentation of macrophages

The M1- and M2-like macrophages were prepared by stimulating bone marrow-derived macrophages (BMDMs) according to our previously reported protocols^24^. For DQ-OVA degradation assay, M1- and M2-like macrophages were plated on glass bottom dishes and treated with various SPEN or SPIN (equivalent to 1 μM IMDQ) with or without irradiation for 24 h. Then the cells were incubated with DQ-OVA (10 μg mL^-1^, Invitrogen) for 15 min, washed, and further incubated at 37 °C for another 15 min. Finally, the cellular fluorescence of DQ-OVA was observed by CLSM. To investigate the antigen presentation efficacy, M2-like macrophages were treated with PEPA or SPEN-stressed and CT26-OVA-derived pyroptosomes for 24 h. Cells were collected, and immunostained with PE anti-mouse H-2K^d^ or PE anti-mouse SIINFEKL-H-2K^b^, respectively. The antigen presentation efficacy of macrophages was then quantified by flow cytometry. The presentation of OVA antigen on cell membrane was further visualized by CLSM after the treated macrophages were labelled with WGA-FITC.

### Multicolor fluorescence immunohistochemistry and analysis

The immune microenvironment in tumours and LNs was visualized by the tyramide signal amplification (TSA) staining using an Opal 7-Color Manual IHC Kit (Akoya Biosciences) according to the manufacturer’s instructions. First, the formalin-fixed and paraffin-embedded tumour and LN tissues were sectioned into 5 μm slides. The slides were deparaffinized with xylenes, rehydrated in decreasing graded alcohol, and antigen retrieved with 1× Tris-EDTA (pH 9.0) in a microwave oven. Then the slides were sequentially blocked with 3% H_2_O_2_ and blocking buffer at room temperature, followed by incubation with primary antibody. After incubation at 4 °C overnight, the sections were washed with PBS for three times, and incubated with HRP-conjugated secondary antibody for 1 h at room temperature. The sections were labelled with Opal probe (1:400) for 10 min (r.t.), and processed to the second round of staining after removal of the Opal probe with 1× Tris-EDTA (pH 9.0) in a microwave oven. The staining sequences for tumour and LN slides were CD8α/CD4/NKp46/Foxp3/CD206/iNOS and CD8α/CD4/CD11c/F4/80, respectively. Finally, the slides were labelled with DAPI, and mounted with prolong diamond antifade mountant. The multispectral images were obtained with a Vectra Polaris system and the quantification analysis was performed with the InForm software.

### In vivo anti-tumour immunotherapy

A bilateral CT26 tumour-bearing mouse model was established by inoculation of 1×10^6^ and 2×10^5^ CT26 cells in the right and left flanks of mice, respectively. When the volume of right tumour reached ∼50 mm^3^, mice were randomly divided into five groups (n = 9-10) and i.v. injected with saline, PEPA, SPIN, and SPEN (equivalent to 1 mg kg^-1^ PPa and 0.4 mg kg^-1^ IMDQ) on day 6 and day 11, respectively. EE stress was elicited by 660 nm-irradiation (240 J cm^−2^) on the right tumours at 3 h post-injection. The tumour size and body weight were monitored every other day, and the survival curve of mice were recorded according to Kaplan-Meier analysis. Mice were euthanized when the tumours reached ∼1,500 mm^3^ in size. For the therapeutic efficiency on large tumours, B16 tumour-bearing mice with tuomur volume of ∼400 mm^3^ was established and treated with the same procedure as described above. For metastasis inhibition study, the therapy of SPEN_EE_ was repeated on 4T1-luc tumour-bearing mice. Bioluminescence imaging with IVIS system was applied to monitor the tumour growth and lung metastasis. On day 35, lung tissues were dissected for luminescence imaging and H&E staining. For tumour rechallenge study, 5×10^5^ MC38 cells were inoculated in SPEN_EE_-treated MC38 tumour-bearing C57BL/6 mice on day 30, 60, and 90, respectively, to investigate the long-term anti-tumour immunity of SPEN_EE_.

### Depletion of macrophages, CD8^+^ T cells, and NK cells in vivo

The CT26 tumour-bearing mice were intraperitoneally injected with anti-mouse CD8 (400 μg per mice for twice weekly), anti-mouse CSF1R (300 μg per mouse, every five days for twice), and anti-ASGM1 (100 μg per mice for twice weekly) before SPEN vaccine therapy to deplete the CD8^+^ T cells, macrophages, or NK, respectively. Depletion of lymphocytes was confirmed by flow cytometry of peripheral blood mononuclear cells.

### Combination immunotherapy of SPEN with αPD-1

PANC02 tumour-bearing C57BL/6 mice were randomly divided into four groups (*n* = 11-12) and treated with saline, αPD-1, SPEN_EE_, and αPD-1 + SPEN_EE_, respectively. On day 8 and day 15, mice were administrated with SPEN (equivalent to 1 mg kg^-1^ PPa and 0.4 mg kg^-1^ IMDQ) via i.v. injection, and the tumors were irradiated with 660 nm laser (240 J cm^−2^) at 3 h post-injection. Mouse αPD-1 (100 μg per mouse, BioXcell) was administered intraperitoneally on day 8, 11, 14, and 17. On day 20, the immunized mice were rechallenged with 5×10^5^ PANC02 cells. The tumour volumes and survival curves were monitored.

### Long-term memory T lymphocytes activation

MC38 tumour-bearing mice were treated with PEPA or SPEN-mediated EE stress once a week for twice. The spleens were excised one months after the second treatment to prepare single cell suspension, and then stained with FITC anti-mouse CD3, PE anti-mouse CD4, APC anti-mouse CD8α, PE/Cy7 anti-mouse CD44, and Pacific Blue anti-mouse CD62L. The CD44^high^CD62L^low^ effector memory T cells and CD44^high^CD62L^high^ central memory T cells were analyzed by flow cytometer.

### Statistics and reproducibility

Data are presented as mean ± s.d. for in vitro and in vivo assays. Statistical analyses were performed using Origin (Origin 2018). An independent-sample t-test or one-way (or two-way) ANOVA followed by Bonferroni’s multiple comparisons test were conducted for analysis of significant differences depending on the number and distribution of treatment groups. Welch’s correction or non-parametric test was applied for the groups without equal s.d. A log-rank test was applied for comparison of survival curves; P<0.05 was considered significant. Each experiment, including transmission electron microscopy, confocal imaging, cell morphology assay and western blot, was independently repeated at least thrice with similar results. The in vivo imaging, immunofluorescence staining of tissues and H&E staining were repeated three times using biologically independent mice with similar results.

### Reporting Summary

Further information on research design is available in the Nature Research Reporting Summary linked to this article.

## Data availability

The main data that support the findings of this study are available within the paper and its Supplementary Information. All raw and relevant data during the study are available for research purposes from the corresponding authors upon reasonable request.

## Acknowledgements

This work was supported by National Key Research and Development Program of China (2021YFA201200 to Y.G.W., 2024YFB3814600 to B.L.C.), National Natural Science Foundation of China (NSFC) Grants (82225044 to Y.G.W., 82373804 and 82450112 to B.L.C.), and Beijing Natural Science Foundation (Z240028 to Y.G.W.). We would like to thank the State Key Laboratory of Natural and Biomimetic Drugs, Peking University Biological Imaging and Flow Cytometry Core Facilities for flow cytometry, confocal, animal, and tissue imaging services.

## Author contributions

B.L.C. and Y.G.W. are responsible for all phases of the research. F.J.W. and L.T.W. helped with fabrication and characterization of the EE-stressed nanophotosensitizer; X.Q.P., J.X.L., and Y.Yang helped with synthesis and characterization of copolymers; H.M.X., Y.Q.W., and M.M.T. participated in the evaluation of immune response; Y.Yan and M.M.Q. helped with in vivo anti-tumour studies; J.N.R. assisted in western blot; W.C. performed HPLC analysis; L.J.Z. conducted the shotgun proteomics analysis; Q.Z. provided conceptual suggestions; Y.G.W. and B.L.C. provided constructive guidance for the overall design of the project; B.L.C. wrote the manuscript; Y.G.W. and B.L.C. revised the paper; All the authors discussed the results and assisted in the preparation of the manuscript.

## Competing interests

The authors declare no competing financial interests.

---

**Extended Figure 1.**
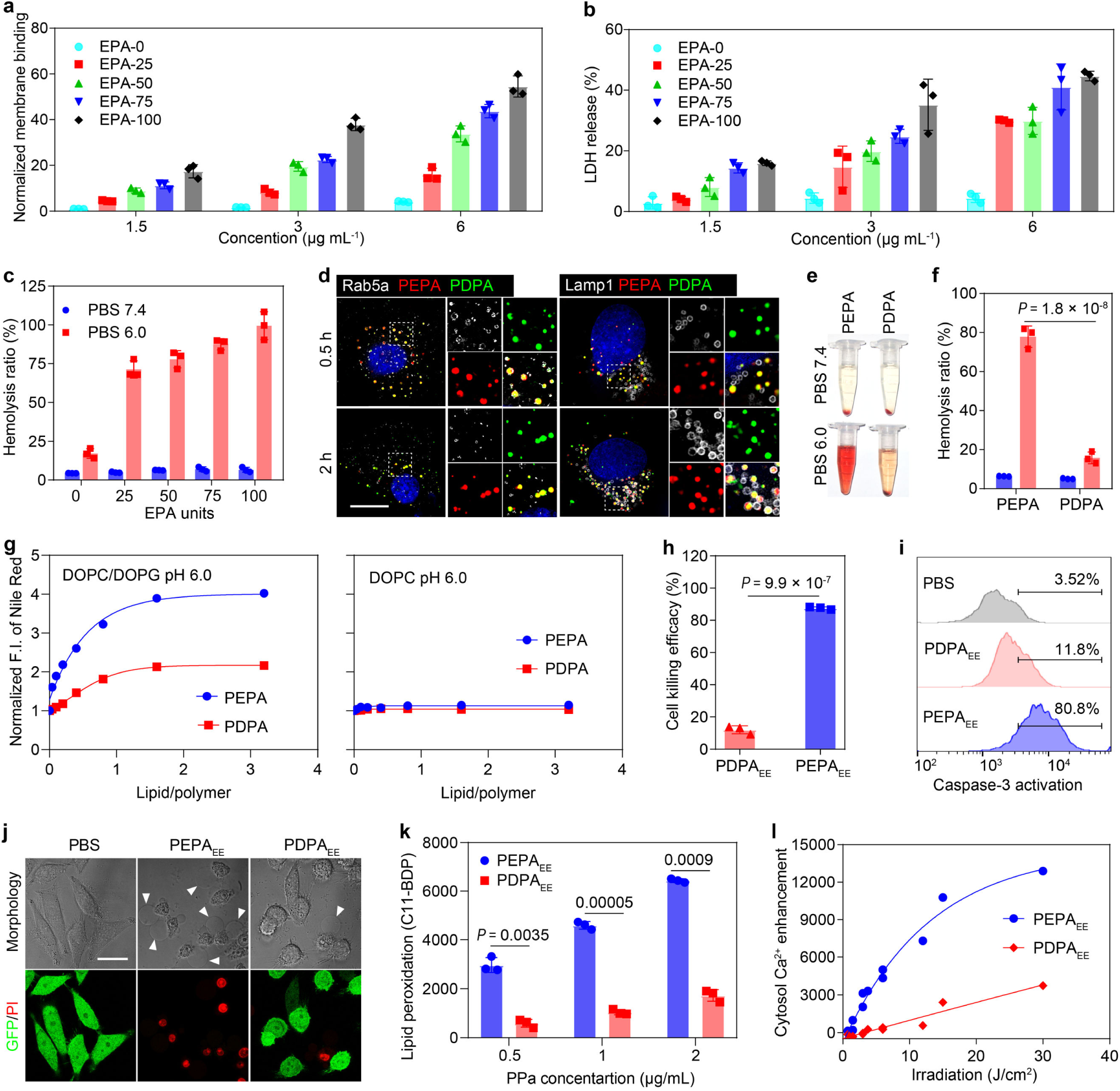
Nanophotosensitizers containing EPA monomers exhibit superior membrane affinity and early endosome stress-mediated cytotoxicity. **(a)** Cell membrane binding capacity of nanophotosensitizers with various EPA monomers on A549 cells at pH 6.0. Fluorescence signal was measured by flow cytometry and normalized to EPA-0 group. **(b)** Oxidative stress-induced LDH release of nanophotosensitizers with various EPA monomers on A549 cells after treatment of membrane binding and 660 nm irradiation (6 J cm^−2^). **(c)** Quantification of erythrocytes hemolysis induced by the nanophotosensitizer-mediated oxidative stress when binding with membrane at pH 7.4 or pH 6.0. **(d)** Confocal images of cellular transport and endocytic organelle colocalization analysis of PEPA and PDPA at 0.5 h and 2 h after internalization. Scale bar = 5 μm. **(e, f)** The membrane-binding of PEPA and PDPA evokes erythrocytes hemolysis with 660 nm irradiation. **(g)** Interaction of Nile red-labelled copolymers with artificial lipid membrane at pH 6.0 measured by fluorimeter. **(h)** Photocytotoxicity of PEPA_EE_ and PDPA_EE_ nanophotosensitizer on A549 cells. **(i)** Intracellular caspase-3 activity of A549 cells measured by flow cytometry with Fluorometric Assay Kit. **(j)** The cell death morphology and PI staining of A549-GFP cells treated with PEPA or PDPA-mediated EE stress. Scale bar = 10 μm. **(k)** Quantification of EE stress-evoked lipid peroxidation detected with oxidation-sensitive sensor C11-BODIPY^581/591^ probe. **(l)** Cytosolic calcium release elicited by the EE stress of PEPA or PDPA. All data are presented as mean ± s.d. (*n* = 3 biologically independent experiments).

**Extended Figure 2.**
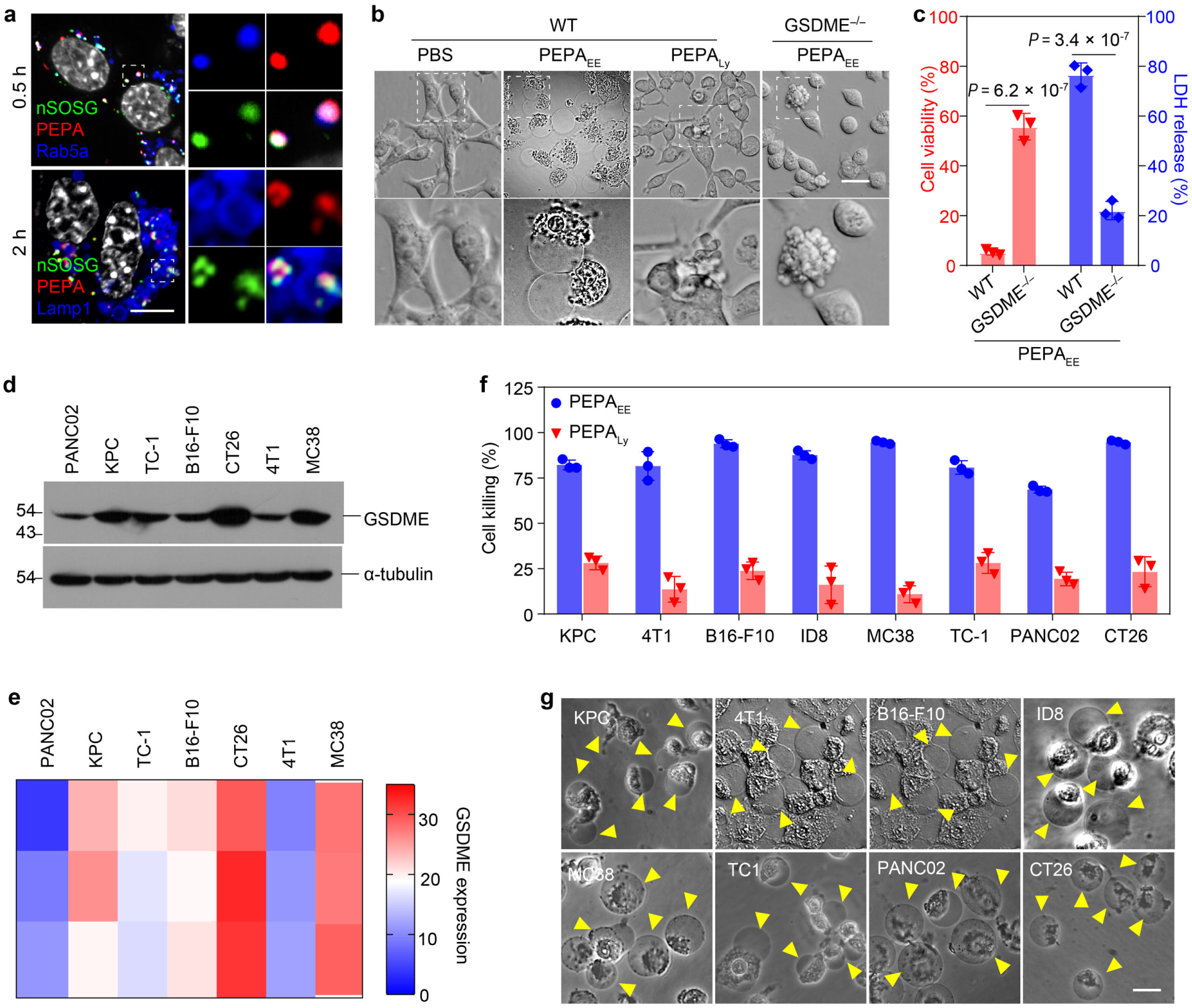
Early endosome stress of PEPA nanophotosensitizer evokes GSDME-mediated pyroptosis in murine tumor cells. **(a)** Confocal images of intracellular singlet oxygen generation measured by SOSG nanoprobe (green) and colocalization with PEPA and endocytic organelles at 0.5 h and 2 h post-internalization. Scale bar = 10 μm. **(b)** Representative phase-contrast images of cell death morphology induced by PEPA_EE_ or PEPA_Ly_ on CT26 and CT26-GSDME^−/−^ cells. Scale bar = 20 μm. **(c)** MTT assay and LDH release assay of PEPA_EE_ stress on CT26 and CT26-GSDME^−/−^ cells. GSDME knockout rescued EE stress-induced pyroptosis. Data are presented as mean ± s.d. (*n* = 3 biologically independent experiments). **(d)** Immunoblots of endogenous GSDME expression in different murine cancer cell lines. **(e)** Heatmap of GSDME expression quantified from panel d. **(f)** Photocytotoxicity of PEPA_EE_ or PEPA_Ly_ on different murine cancer cell lines. Data are presented as mean ± s.d. (*n* = 3 biologically independent experiments). **(g)** Phase-contrast imaging of cell death morphology on different murine cancer cells treated with PEPA_6.5_-mediated EE stress. Scale bar = 20 μm.

**Extended Figure 3.**
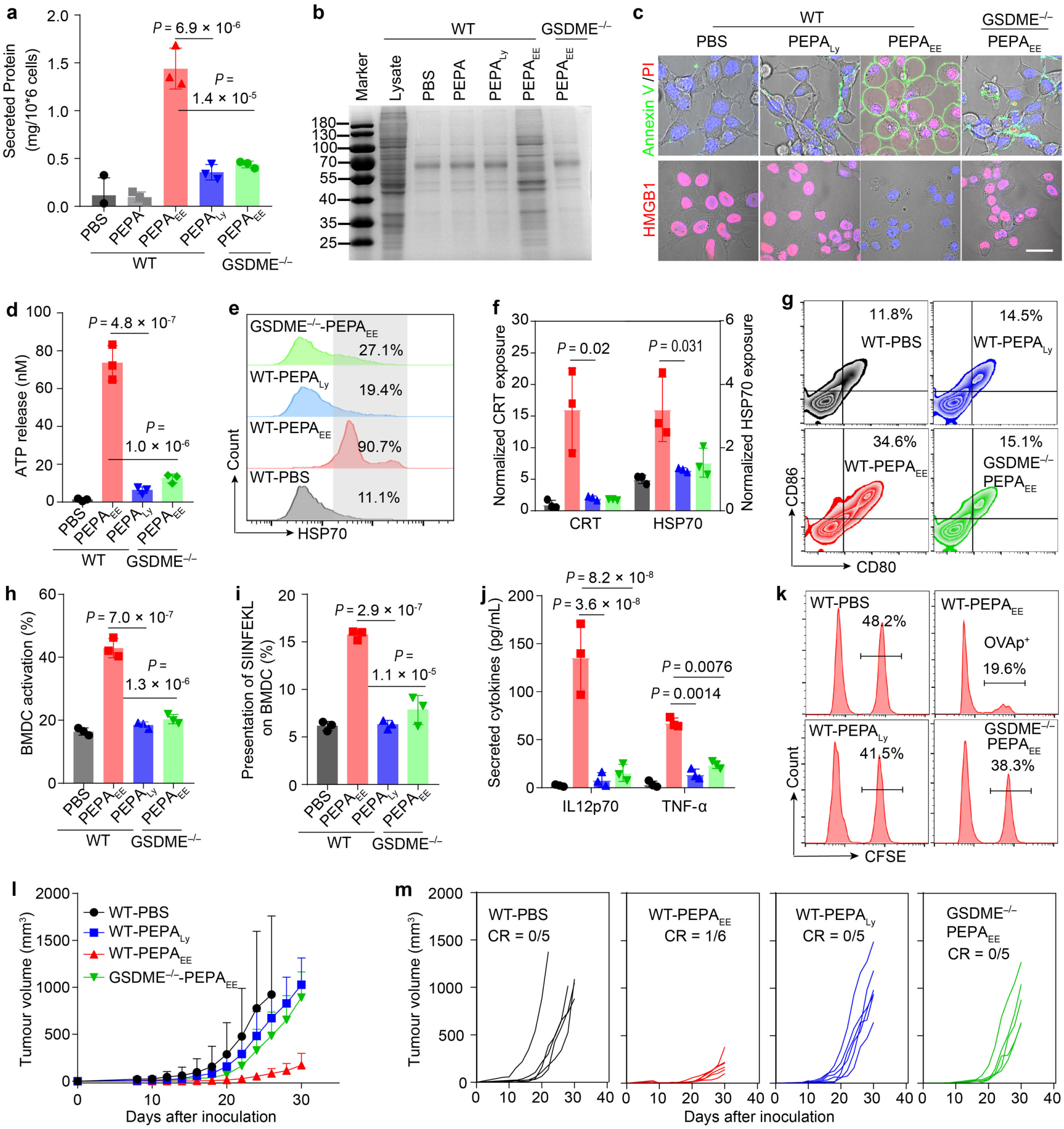
EE stress-evoked pyroptosis enhances the ICD effect of cancer cells in vitro and in vivo. (a, **b)** PEPA-mediated EE stress facilitates intracellular protein release on CT26 cells as evaluated by BCA assay **(a)** and SDS-PAGE **(b)**. *n* = 3 biologically independent experiments. **(c)** Confocal images of annexin V-FITC/PI and HMGB1 staining of CT26 cells at 24 h post-treatment with PEPA_EE_ or PEPA_Ly_. Scale bar = 20 μm. **(d)** ATP release level, and **(e, f)** HSP70/CRT exposure of CT26 cells at 24 h post-treatment with PEPA_EE_ or PEPA_Ly_. *n* = 3 biologically independent experiments. **(g-j)** BMDC activation after co-culture with PEPA_EE_ or PEPA_Ly_ treated CT26-OVA cells. *n* = 3 biologically independent experiments. Representative contour plots of CD80^+^CD86^+^ cells among CD11c^+^ BMDC (g). Percentage of CD80^+^CD86^+^ BMDC (h). Percentage of BMDCs with presentation of OVA antigen **(i)**. Secretion of IL-12 and TNF-α measured by ELISA assay **(j)**. **(k)** Representative results of specific cell killing study in CT26-OVA tumour-bearing mice after different treatments. **(l)** Average tumour growth curves and **(m)** individual tumour growth kinetics of CT26-OVA tumour-bearing mice treated with PEPA_EE_ or PEPA_Ly_. *n* = 5 biologically independent mice. All data are shown as mean ± s.d.

**Extended Figure 4.**
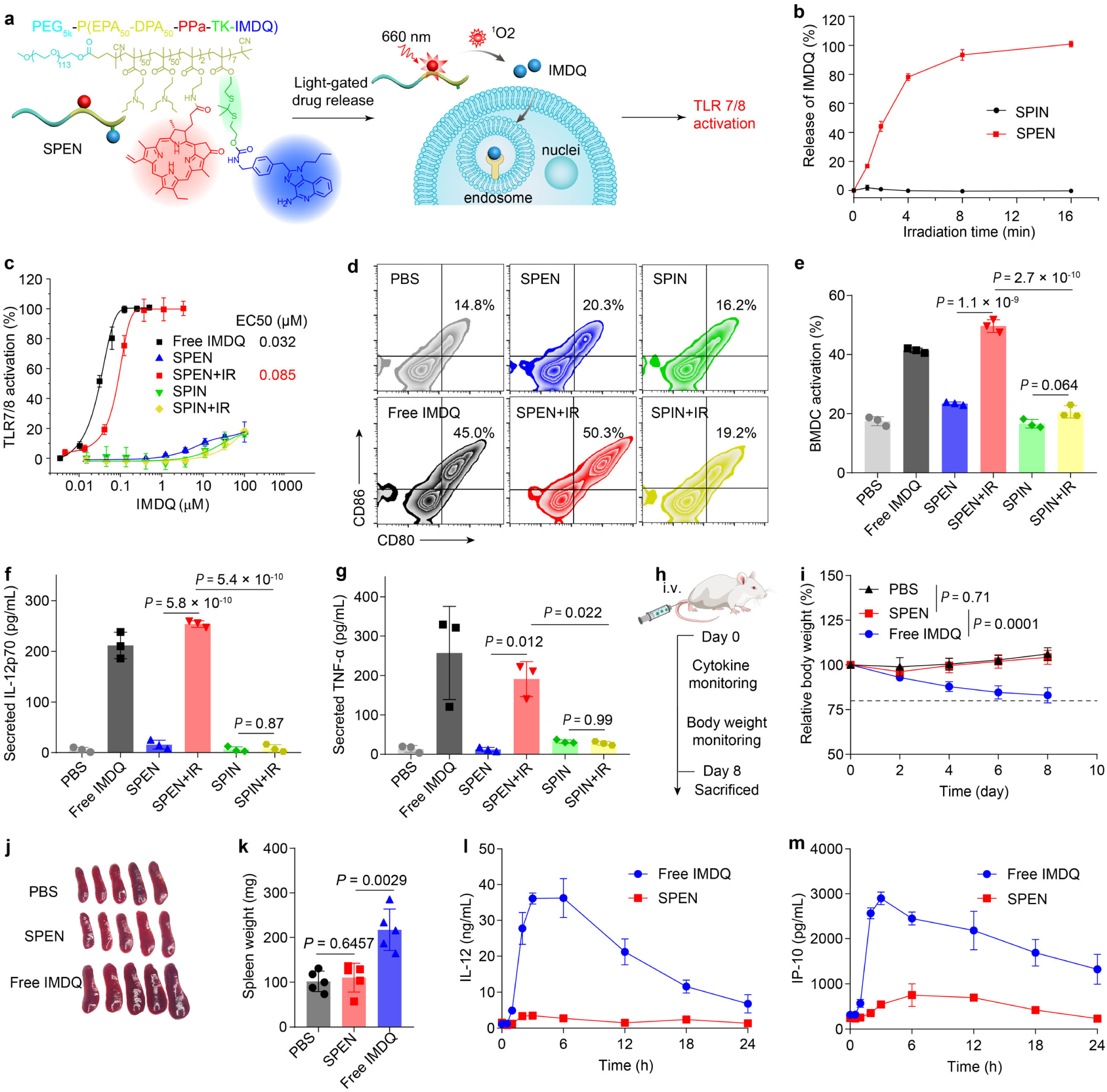
SPEN exhibits safe and effective TLR activation. **(a)** Chemical structure of the pH and light-gated polymer-agonist conjugate and schematic of light-triggered TLR activation. **(b)** 660 nm laser-induced release profiles of SPEN and SPIN quantified by HPLC (*n* = 3 biologically independent experiments). **(c)** TLR agonistic activity of various IMDQ treatments with or without 660 nm irradiation via a TLR reporter cell assay (*n* = 6 biologically independent experiments). **(d-g)** BMDC activation after treatment of free IMDQ, SPEN, or SPIN with or without 660 nm irradiation. n = 3 biologically independent experiments. Representative contour plots of CD80^+^CD86^+^ cells among CD11c^+^ BMDC **(d)**. Percentage of CD80^+^CD86^+^ BMDC **(e)**. Secretion of IL-12 **(f)** and TNF-α **(g)** measured by ELISA assay. **(h-m)** In vivo biosafety and systemic immunotoxicity of SPEN (*n* = 5 biologically independent mice). Mice were administrated with free IMDQ or SPEN via tail vein injection, and the body weight, cytokines in blood, and spleens were evaluated **(h)**. Mice body weight change after different treatments **(i)**. Photographs **(j)** and weight **(k)** of spleens excised at day 8. The profiles of IL-12 **(l)** and IP-10 **(m)** level in blood. All data are shown as mean ± s.d.

**Extended Figure 5.**
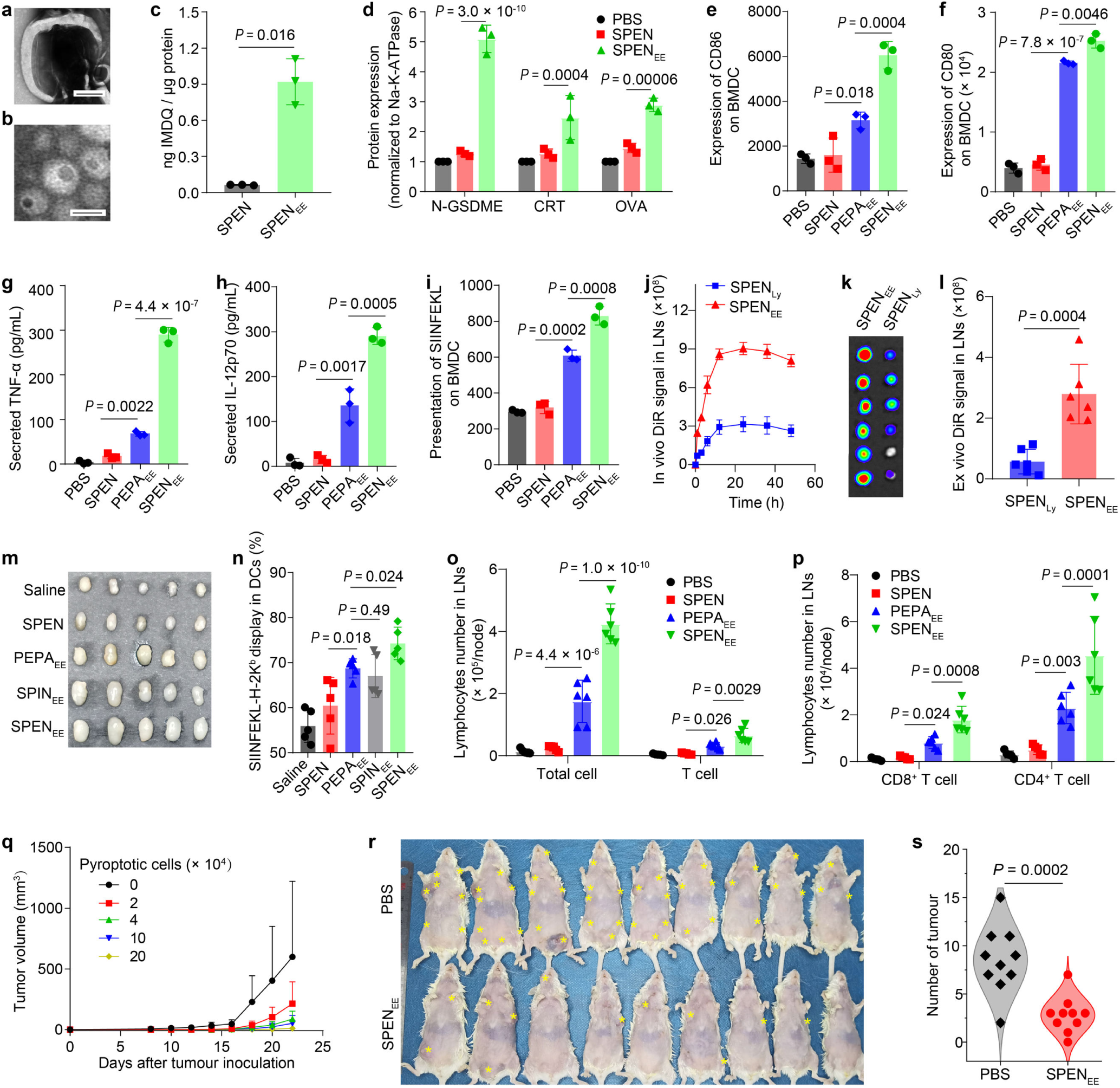
In vitro and in vivo cancer vaccination capacity of EE stress-derived pyroptosomes. **(a)** Morphology of EE stress-derived pyroptosome characterized by TEM. Scale bar = 500 nm. **(b)** Morphology of Au nanoparticle-encapsulated SPEN characterized by TEM. Scale bar = 20 nm. (**c**) Enrichment of GSDME-N, calreticulin (CRT), and ovalbumin (OVA) in the cell membrane or pyroptosomes derived from CT26-OVA (normalized to Na-K-ATPase, *n* = 3 biologically independent experiments). **(d)** STING activation efficacy of pyroptosomes via Raw-Blue-ISG reporter cell assay (*n* = 3 biologically independent experiments). **(e-i)** BMDC activation after treatment with pyroptosomes elicited by EE stress of various nanoadjuvants (n = 3 biologically independent experiments). Quantitative expression of CD86 **(e)** and CD80 **(f)** on BMDCs. Secretion of TNF-α **(g)** and IL-12 **(h)** measured by ELISA assay. Presentation of OVA antigen **(i). (j-l)** In vivo lymph node (LN) draining of DiR-labelled pyroptosomes (*n* = 3 biologically independent mice). Quantitative DiR signal profiles in popliteal LNs **(j)**. Ex vivo fluorescence imaging **(k)** and quantification **(l)** of excised LNs. **(m-p)** Activation of LNs in mice immunized with pyroptosomes (*n* = 5 biologically independent mice). Photographs of excised LNs **(m).** Presentation of OVA antigen on DCs in LNs **(n)**. The cell counts of total lymphocytes and T cells in LNs **(o)**. The cell counts of CD8^+^ T cells and CD4^+^ T cells in LNs **(p)**. **(q)** Tumour growth profiles of CT26 tumour-bearing mice immunized with pyroptosomes from 0, 2, 4, 10 or 20 × 10^4^ pyroptotic cells. **(r)** Photographs and **(s)** numbers of spontaneous tumours on MMTV-PyMT/FVB mice immunized with personalized pyroptotic cancer vaccine (n = 10 biologically independent mice; two-sided Student’s t-test with Welch’s correction). All data are shown as mean ± s.d.

**Extended Figure 6.**
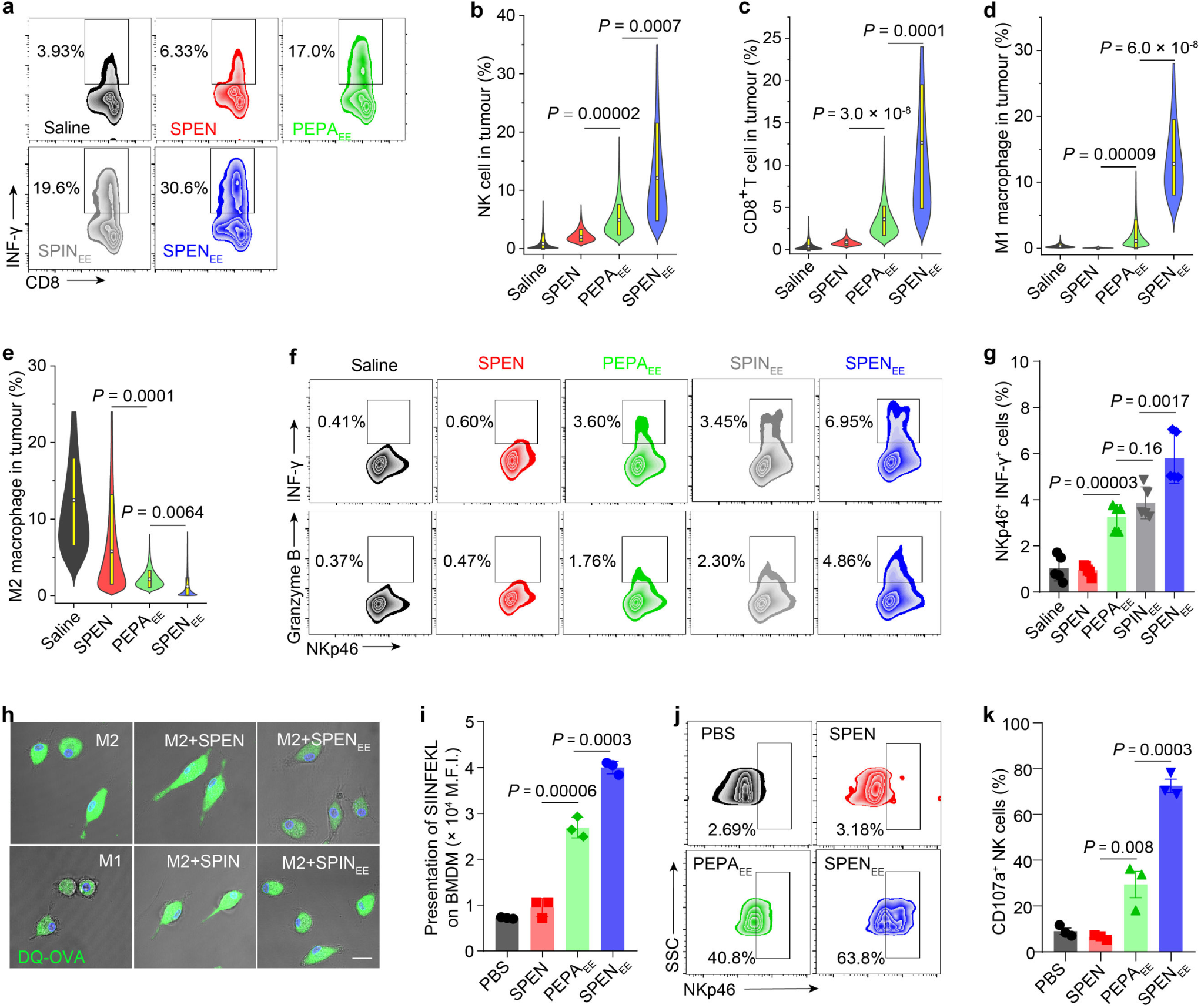
Tumour immune microenvironment priming of SPEN-mediated EE stress. **(a)** Representative flow cytometry plots of INF-γ^+^ cytotoxic T cells among CD8^+^ T cells in immunized mice (*n* = 5 biologically independent mice). **(b-e)** The percentage of NK cells **(b)**, CD8^+^ T cells **(c)**, M1-type macrophages **(d)**, and M2-type macrophages **(e)** in tumour tissues quantified from multiplex immunohistochemistry staining of tumour slides (*n* = 20-30 biologically independent regions). **(f)** Representative flow cytometry plots of INF-γ^+^ NK cells and granzyme B^+^ NK cells among NKp46^+^ cells in immunized mice. **(g)** The percentage of INF-γ^+^ cytotoxic NK cells among NKp46^+^ cells in immunized mice (*n* = 5 biologically independent mice). **(h)** Representative confocal images of DQ-ovalbumin (green) degradation assays in BMDMs pretreated with various pyroptosomes. Scale bar = 20 μm. **(i)** Presentation of OVA antigen on cell membrane of BMDM after treatment with various pyroptosomes (*n* = 3 biologically independent experiment). **(j)** Representative flow cytometry plots of NK activation by various pyroptosomes as indicated by membrane expression of NKp46. **(k)** Quantification of CD107a^+^ cells among NKp46^+^ NK cells (*n* = 3 biologically independent experiment). All data are shown as mean ± s.d. Statistical significance was analyzed by one-way ANOVA followed by Dunnett’s multiple comparisons test.

**Extended Figure 7.**
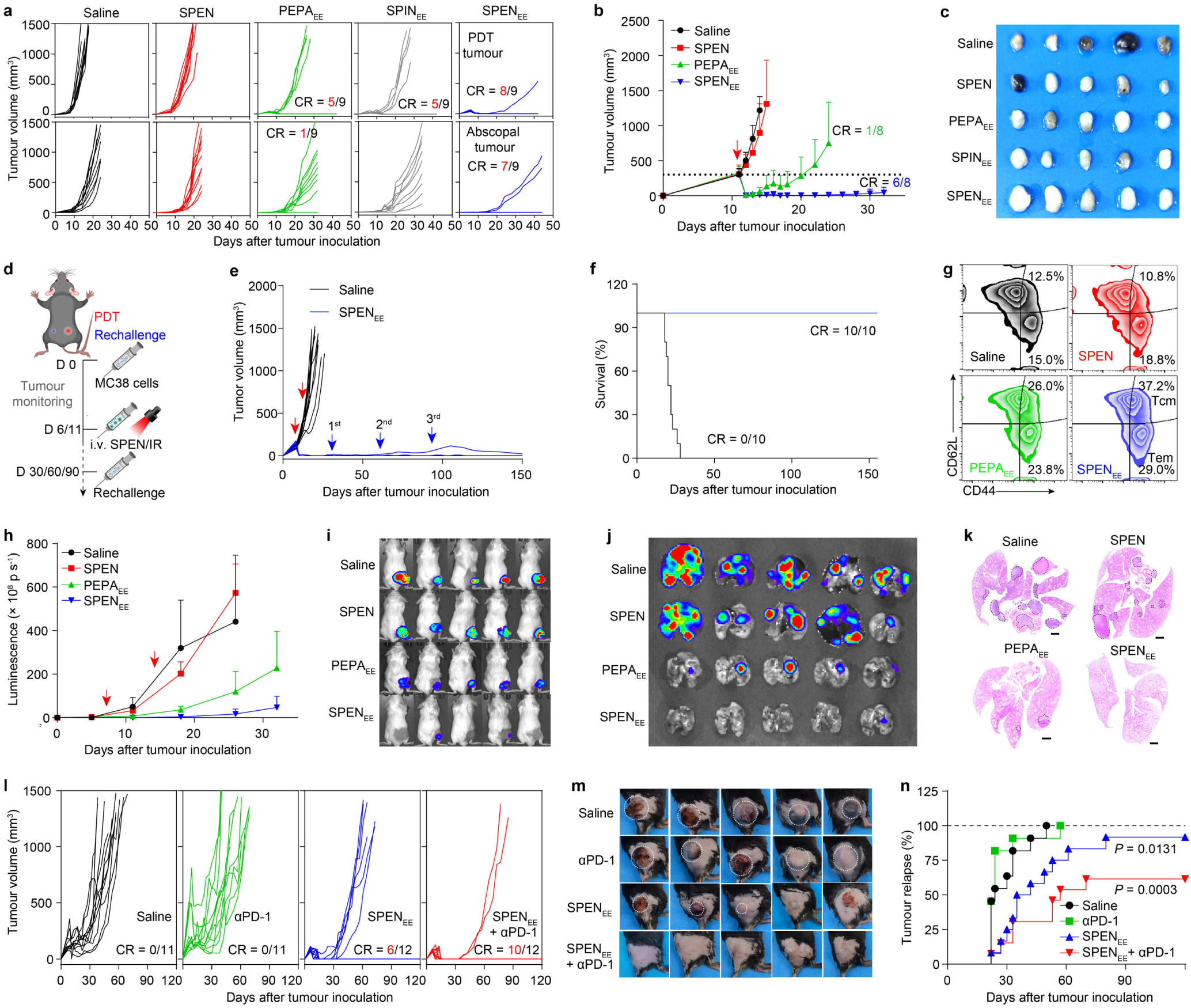
In vivo cancer immunotherapy of SPEN-mediated EE stress. **(a)** Individual tumour growth curves of CT26 tumour-bearing mice treated with SPEN_EE_-mediated in situ vaccination (*n* = 10 mice for saline and SPEN groups; *n* = 9 mice for PEPA_EE_, SPIN_EE_, and SPEN_EE_ groups). **(b)** Tumour growth profiles and **(c)** excised metastatic LNs of large B16 tumour bearing mice with treatment of SPEN_EE_-mediated in situ vaccination. *n* = 7 mice for saline and SPEN groups; *n* = 8 mice for PEPA_EE_ and SPEN_EE_ groups. **(d-g)** Long-lasting cancer prevention capacity of SPEN_EE_ on MC38 tumour-bearing mice. Schematic illustration of rechallenge study **(d)**. Individual tumour growth curves **(e)** and animal survival curves **(f)** of MC38 tumour-bearing mice treated with SPEN_EE_-mediated in situ vaccination (*n* = 10 biologically independent mice). Representative scatterplots of CD44^high^ CD62L^low^ T cell and CD44^high^ CD62L^high^ T cell subsets among CD4^+^ T lymphocytes **(g)**. **(h-k)** Immunotherapy and metastasis inhibition efficacy on 4T1-luc tumour-bearing mice (*n* = 5 biologically independent mice). Tumour growth curves quantified by in vivo luminescence imaging **(h)**. The luminescence imaging of 4T1-luc tumour bearing mice at day 32 **(i)**. The photographs of luminescence imaging **(j)** and H&E staining **(k)** of excised lungs. **(l)** Individual tumour growth curves, **(m)** representative photographs of tumour-bearing mice at day 60, and **(n)** tumour relapse curves of the combination immunotherapy of SPEN_EE_ with αPD-1 on the immunosuppressive PANC02 tumour-bearing mice (*n* = 11 mice for saline and αPD-1 groups; *n* = 12 mice for SPEN_EE_ and SPEN_EE_ + αPD-1 groups; log-rank test; *P* = 0.0131 for αPD-1 versus SPEN_EE_, *P* = 0.0003 for SPEN_EE_ versus SPEN_EE_ + αPD-1). All data are shown as mean ± s.d.

## Notes

### Competing Interest Statement

The authors have declared no competing interest.

