## Supplementary Information for "In situ generated vaccine-like pyroptosome for personalized cancer immunotherapy"

[1]State Key Laboratory of Natural and Biomimetic Drugs, School of Pharmaceutical Sciences, Peking University, Beijing 100191, China. [2]Beijing Key Laboratory of Molecular Pharmaceutics and New Drug Delivery System, School of Pharmaceutical Sciences, Peking University, Beijing 100191, China. [3]College of New Materials and Chemical Engineering, Beijing Institute of Petrochemical Technology, Beijing, 102617, China. [4]School of Pharmaceutical Sciences, Capital Medical University, Beijing 100069, China. [5]Department of Central Laboratory, Peking University First Hospital, Beijing 100034, China. [6]Center of Medical and Health Analysis, Peking University Health Science Center, Beijing 100191, China. [7]Chemical Biology Center, Peking University, Beijing 100191, China.

* (B.C.); (Q.Z.); (Y.W.)

**Materials and antibodies.** The photosensitizer pyropheophorbide-a (PPa) was purchased from Shanghai Xianhui Pharmaceutical Co., Ltd. The fluorescence probe Cy7.5-NHS ester was purchased from Lumiprobe Company. N-Hydroxysuccinimide (NHS), dicyclohexylcarbodiimide (DCC), MeO-$PEG_{114}$-OH, N,N,N',N",N"-pentamethyldiethylenetriamine (PMDETA), and 3-(4,5-dimethylthiazol-2-yl)-2,5-diphenyltetrazolium bromide (MTT) were all obtained from Sigma-Aldrich. The TLR agonist, imidazoquinoline (IMDQ), was obtained from Nanjing Aikon Chemical Ltd. The monomers 2-(diisopropylamino) ethyl methacrylate (iDPA-MA) and 2-aminoethyl methacrylate (AMA) were purchased from Polyscience Company. The macroinitiator MeO-$PEG_{114}$-Br, monomers 2-(dipropylamino) ethyl methacrylate (nDPA-MA), 2-(dibutylamino) ethanol methacrylate (DBA-MA), and 2-(ethylpropylamino) ethyl methacrylate (EPA-MA) were all synthesized according to our published procedures[1, 2]. 1-palmitoyl-2-oleoyl-*sn*-glycero-3- phosphocholine (POPC) and 1-palmitoyl-2-oleoyl-*sn*-glycero-3-phosphoglycerol (POPG) were purchased from NOF Corporation. Other solvents and reagents were received from Sigma-Aldrich or Fisher Scientific Inc.

Anti-Ki-67 (ab15580), HMGB1 (ab79823), GSDME (ab215191) antibodies, CD8 (ab217344), NKp46 (ab233558), Foxp3 (ab215206), and CD4 (ab183685) were purchased from Abcam. Antibody for caspase-3 (#9662), CD11c (#97585), and F4/80 (#70076) were obtained from Cell Signaling Technology. Anti-tubulin antibody (T5168) was purchased from Sigma-Aldrich. $OVA_{257-264}$ peptide was purchased from Sangon Biotech. Fluorescence labeled anti-mouse monoclonal antibodies for flow cytometry analysis were purchased from Biolegend Company. ELISA kits for the detection of IP-10 and IL-12 were purchased from Peprotech.

**Synthesis of IMDQ methacrylate monomers.** The ROS-inactivated HHMA-IMDQ (**Fig.S4**) was synthesized as previously reported[3]. The ROS-activatable MA-TK-IMDQ was synthesized according to the procedures in **Fig.S5**.

**Synthesis of TK-COOH:** A mixture of anhydrous mercaptoacetic acid (4.20 g, 50.0 mmol) and anhydrous acetone (5.80 g, 100.0 mmol) were saturated with boron trifluoride and stirred at room temperature (r.t.) for 6 h. The reaction mixture was crystallized with ice-salt bath, washed with hexane and cold water, and lyophilized to obtain TK-COOH (85.8% yield). ESI-MS of $C_7H_{12}O_4S_2$ was shown in **Fig.S6a**. $^1$H NMR (400 MHz, $CDCl_3$, shown in **Fig.S6b**): δ 12.61 (s, 2H), 3.36 (s, 4H), 1.54 (s, 6H).

**Synthesis of TK-OH:** 4.00 g (17.86 mmol) of TK-COOH was dissolved in 50 mL of anhydrous tetrahydrofuran (THF), and 44.64 mL (2.4M in THF, 107.13 mmol) of $LiAlH_4$ was added to the solution dropwise in ice bath under stirring. After reaction for 8 h (r.t.), the crude product was purified via column chromatography (DCM/EA = 50/50) to obtain yellow oily TK-OH (61.1% yield). $C_7H_{16}O_2S_2$, ESI-MS m/z: 195.0513 $[M-H]^-$, 241.0567 $[M+HCOO]^-$ (**Fig.S7a**). $^1$H NMR (400 MHz, DMSO-$d_6$, shown in **Fig.S7b**): δ 4.82 (t, 2H), 3.52 (q, 4H), 2.65 (t, 4H), 1.52 (s, 6H).

**Synthesis of MA-TK-OH:** A mixture of TK-OH (1.55 g, 7.91 mmol) and TEA (1.20 g, 11.86 mmol) were dissolved in 60 mL of anhydrous DCM, and then 0.83 g of methacryloyl chloride (7.91 mmol in DCM) was added dropwise under stirring at 0 °C. The reaction mixture was stirred for 12 h (r.t.), and then purified via column chromatography (DCM/EA = 90/10). Finally, a colorless liquid was obtained by drying in vacuo overnight (35.9% yield). $C_{11}H_{20}O_3S_2$, ESI-MS m/z: 287.0740 $[M+Na]^+$ (**Fig.S8a**). $^1$H NMR (400 MHz, DMSO-$d_6$, shown in **Fig.S8b**): δ 6.04 (s, 1H), 5.70 (t, 1H), 4.82 (t, 1H), 4.24 (t, 2H), 3.52 (q, 2H), 2.87 (t, 2H), 2.66 (t, 2H), 1.88 (s, 3H), 1.55 (s, 6H).

**Synthesis of MA-TK-PNP:** MA-TK-OH (1.47 g, 5.57 mmol) and TEA (0.85 g, 8.35 mmol) were dissolved in 25 mL of anhydrous DCM. Then 1.23 g of *p*-nitrophenyl chloroformate (6.12 mmol) was dissolved in 10 mL of anhydrous DCM, and dropwise added into the mixture in ice bath under stirring. Following 5 h of stirring (r.t.), the crude product was purified via column chromatography (PE/EA = 90/10). A white powder product was obtained by drying in vacuo overnight (89.7% yield). $C_{18}H_{23}N_1O_7S_2$, ESI-MS m/z: 447.1298 $[M+NH4]^+$ (**Fig.S9a**). $^1$H NMR (400 MHz, DMSO-$d_6$, shown in **Fig.S9b**): δ 8.32 (d, 2H), 7.56 (d, 2H), 6.03 (s, 1H), 5.69 (t, 1H), 4.39 (t, 2H), 4.26 (t, 2H), 2.97 (t, 2H), 2.90 (t, 2H), 1.87 (s, 3H), 1.59 (s, 6H).

**Synthesis of MA-TK-IMDQ:** A mixture of IMDQ (0.30 g, 0.70 mmol) and TEA (0.28 g, 2.78 mmol) were dissolved in 10 mL of anhydrous DMSO, and then 0.30 g of MA-TK-PNP (0.70 mmol) in 5 mL of anhydrous DMSO was dropwise added into the mixture. The above mixture was kept at room temperature under a nitrogen atmosphere overnight. After purification via column chromatography (DCM/MeOH = 95/5), a white solid product was obtained by drying in vacuo overnight (59.8% yield). $C_{34}H_{43}N_5O_4S_2$, ESI-MS m/z: 650.2901 $[M+H]^+$ (**Fig.S10a**). $^1$H NMR (400 MHz, DMSO-$d_6$, shown in **Fig.S10b**): δ 7.80 (d, 1H), 7.70 (t, 1H), 7.60 (d, 1H), 7.36 (t, 1H), 7.20 (d, 2H), 7.07 (t, 1H), 6.99 (d, 2H), 6.84 (s, 2H), 6.02 (s, 1H), 5.84 (s, 2H), 5.67 (s, 1H), 4.22 (t, 2H), 4.09 (m, 4H), 2.86 (m, 4H), 2.76 (t, 2H), 1.86 (s, 3H), 1.70 (m, 2H), 1.53 (s, 6H), 1.37 (m, 2H), 0.86 (t, 3H).

**Synthesis and characterization of polymer-drug conjugates.** The IMDQ-conjugate copolymer-drug conjugates PEG-*b*-P($EPA_{50}$-*r*-$DPA_{50}$-*r*-IMDQ-$AMA_3$) and PEG-*b*-P($EPA_{50}$-*r*-$DPA_{50}$-*r*-TK-IMDQ-$AMA_3$) were synthesized via reversible addition-fragmentation chain transfer (RAFT) polymerization method[4] as shown in **Fig.S11**. Briefly, take PEG-*b*-P($EPA_{50}$-*r*-$DPA_{50}$-*r*-TK-IMDQ-$AMA_3$) as an example, $PEG_{5k}$-CTA (0.5 g, 0.1 mmol), EPA-MA (995 mg, 5.0 mmol), DPA-MA (1065 mg, 5.0 mmol), AMA (50 mg, 0.3 mmol), MA-TK-IMDQ (325 mg, 0.5 mmol), and AIBN (4.1 mg, 25 μmol) were mixed into a polymerization tube with anhydrous DMF, followed with three cycles of freeze-pump-thaw to keep an oxygen-free environment. After polymerization for 48 h at 65 °C, the reaction was successively dialyzed in DMF and Milli-Q water for 48 h. The polymer-drug conjugates were finally obtained by lyophilization, and further characterized by 400 MHz $^1$H-NMR (**Fig.S12, S13**) and gel permeation chromatography in **Supplementary Table 1**.

For photosensitizer conjugation, PPa (10.7 mg, 0.02 mmol), DCC (4.5 mg, 0.022 mmol) and NHS (2.8 mg, 0.024 mmol) were mixed in a reaction bottle, and then dissolved in 0.2 mL anhydrous DMF. The reaction mixture was stirred for 4 h (r.t.), and then PEG-*b*-P($EPA_{50}$-*r*-$DPA_{50}$-*r*-TK-IMDQ-$AMA_3$) copolymers (94 mg, 0.004 mmol) were dissolved in 0.4 mL anhydrous DMF and added to the reaction mixture for another 24 h reaction. Subsequently, the resulting photosensitizer-conjugated copolymers were purified by preparative gel permeation chromatography, lyophilized, and kept at -20 °C for storage.

**Light-activated release of IMDQ.** The 660 nm laser-gated and ROS-induced release profile of IMDQ from SPEN was evaluated by high performance liquid chromatograph (HPLC). The SPIN and SPEN were dispersed into 100 mM PBS buffers with pH of 5.4, respectively, and irradiated with a 660 nm laser (100 mW cm$^{-2}$). At predesignated time-points, the solution was sampled and the released amount of IMDQ was detected by HPLC at a UV wavelength of 322 nm. Chromatographic column: ZORBAX Eclipse Plus C8. Mobile phase: acetonitrile: 0.3% acetic acid = 10:90 - 90:10.

**Membrane binding and rupture assay.** For membrane binding assay, A549 cells were seeded into 24-well plates at a density of 50,000 cells per well and incubated at 37 °C overnight for adherence. Then the culture medium was replaced with fresh medium of pH 5.4 containing nanophotosensitizers with various EPA units, respectively. After cell incubation at 4 °C for 30 min, the cells were trypsinized, harvested, and washed with PBS. The membrane binding of nanoparticles was quantified by flow cytometry (CytoFLEX LX, Beckman, CytExpert 2.3) and IVIS Spectrum imaging ($\lambda_{ex}/\lambda_{em}$: 620 ± 10 nm/ 670 ± 20 nm).

For confocal laser scan microscopy (CLSM) observation, A549 cells were plated on a glass bottom dish and cultured overnight. After the cells were incubated with PEPA or PDPA, 1 µg mL$^{-1}$ of FITC-labeled wheat germ agglutinin (WGA, Molecular Probe) and 5 µg mL$^{-1}$ of Hoechst 33342 (Molecular Probe) were used to indicate the cell membrane and nuclei, respectively. Images were captured with an A1R-Storm confocal microscope ((NIS-Elements AR 4.20.00, Nikon). The DAPI, FITC, and Cy5 filters were used for Hoechst 33342, FITC, and PPa imaging, respectively.

The nanophotosensitizer-treated cells were also irradiated with a 660 nm laser for 1 min (100 mW cm$^{-2}$, 6 J cm$^{-2}$) to investigate the nanoparticle-induce membrane rupture by LDH release assay (Beyotime, China) according to the manufacturer's instructions.

**Erythrocyte lysis assay.** Fresh red blood cells (RBCs) from mouse were isolated with heparin by centrifuging at 4000 rpm for 10 min, and washed with 10 mM PBS for three times. The obtained RBCs were suspended in PBS with pH 7.4 or 5.4 at a density of 1.0 × 10$^{8}$ per milliliter, and incubated with various PEPA nanophotosensitizers (6 µg mL$^{-1}$ PPa) for 0.5 h at 4 °C. Blank PBS of pH 7.4 and distilled water suspended RBCs were included as negative and positive control. After 660 nm laser (100 mW cm$^{-2}$ for 1 min, 6 J cm$^{-2}$), samples were centrifuged and the absorbance of the supernatant at 540 nm was then measured. Hemolysis activity was normalized to the water group as 100% lysis control.

**Interactions between artificial lipid membrane and nanophotosensitizer.** Artificial lipid membranes (ALMs) were fabricated to assess the interactions of nanophotosensitizers with biomembranes by isothermal titration calorimetry (ITC) and Nile red (NR) staining. ALMs formed by DOPC or a mixture of DOPC/DOPG (molar ratio = 5 : 1) were prepared to simulate the cytomembrane or endosome membrane, respectively, as reported previously[5]. Briefly, lipids were dissolved in chloroform-methanol and rotary evaporated to form a lipid film. Then the lipid film was hydrated with PBS solution and extruded 10 times with an Avanti extruder to give large unilamellar vesicles. The ITC assay was performed on a PEAQ ITC (Malvern Instruments) at 25 °C. The DOPC and DOPC/DOPG ALMs were diluted in 100 mM PBS with pH of 7.4 or 6.0 at a lipid concentration of 2.5 mM, respectively. The PEPA or PDPA nanophotosensitizers, at a polymer concentration of 50 μM, were sequentially injected into the ALMs using an injection sequence of 2 μL × 20 with an interval of 2 min. The titration curves were fitted with a model of one binding site using Origin 9.0 (OriginLab).

NR is an environment-sensitive probe exhibiting higher fluorescence in lower polarity, and has been widely applied as a lipid stain to monitor the dynamics in membranes[6]. Therefore, we conjugated NR with PEPA and PDPA to indicate the interaction between nanophotosensitizers with ALMs. The NR-labeled PEPA and PDPA were dispersed in PBS with pH of 6.0 at a polymer concentration of 100 μg $mL^{-1}$, respectively, and increasing ALMs was added to the samples. The fluorescence spectrum of NR ($\lambda_{ex}$ = 575 nm; $\lambda_{em}$ = 590-700 nm) was recorded using a fluorescence spectrophotometer (Hitachi).

**Intracellular singlet oxygen imaging.** The cellular singlet oxygen generation (SOG) of PEPA in distinct endocytic organelles was visualized with the SOSG nanosensor developed by our laboratory[7]. Briefly, A549 cells were co-treated with PEPA and SOSG nanosensor for 0.5 h, and 660 nm irradiation (100 mW $cm^{-2}$ for 1 min) was carried out at 0 h (EE stress) or 2 h (Ly stress) post-incubation to induce SOG in EE or Ly, respectively. The subcellular distribution of singlet oxygen after irradiation was further imaged with confocal microscope. EE and Ly were labelled with RFP for colocalization study. The Hoechst 33342, SOSG, RFP, and PPa were excited at 405, 488, 561, and 633 nm, respectively. The DAPI, FITC, TRITC, and Cy5 filters were used for Hoechst 33342, SOSG, RFP, and PPa imaging, respectively.

**Intracellular lipid oxidation, calcium signal, and caspase-3 activation.** The intracellular lipid peroxide was indicated by a lipid peroxidation probe C11-BODIPY581/591 (Invitrogen). The A549 cells with treatment of PEPA-mediated or PDPA-mediated EE stress were stained with 2 μM C11-BODIPY581/591 for 0.5 h. Cellular lipid peroxide was then quantified by flow cytometry. Fluo3-AM (Beyotime) was used to detect cellular calcium signal according to the manufacturer's instructions. A549 cells were incubated with PEPA or PDPA, followed by staining with 5 μM Fluo3-AM for 15 min. Increasing irradiation dose was carried out immediately after staining, and the cellular Fluo-3 signal was quantified by flow cytometry after various irradiation. GreenNuc™ Caspase-3 Assay Kit (Beyotime) was used to detect caspase-3 activity in living cells. After exposed to $PEPA_{EE}$ or $PDPA_{EE}$ treatments, cells were stained with caspase-3 substrate for 20 min at 37 °C following the manufacturer's procedure. Flow cytometry was applied to quantify caspase-3 activation.

**TLR and STING activation efficacy.** The RAW-Blue and RAW-Blue-ISG reporter cell lines were applied to evaluate the TLR activation[8] and STING activation[9], respectively. To investigate the light-gated TLR activation of SPEN, the RAW-Blue cells were seeded in 96-well plate with 5 $\times 10^4$ cells/well and cultured in DMEM medium supplied with 10% heat-inactivated FBS. After incubation with different IMDQ formulations with or without 660 nm irradiation (100 mW $cm^2$ for 10 min) for 24 hrs, 50 μL of supernatant was collected and mixed with 150 μL Quanti-blue substrate solution (InvivoGen). Then the absorbance at 620 nm was quantified by a Microplate Reader (Multiskan FC, Thermo Fisher Scientific). The TLR activation curves and the EC50 values were fitted and calculated by Origin Pro software. To probe the immune priming effect of pyroptosomes, the RAW-Blue and RAW-Blue-ISG cells were treated with different EE-stressed and CT26-derived pyroptosomes for 24 hrs, respectively. Then the supernatants were collected for Quanti-blue colouring.

**Biosafety and cytokine storm evaluation.** Healthy BALB/c mice were intravenously (i.v.) injected with free IMDQ or SPEN (equivalent to 0.4 mg $kg^{-1}$ IMDQ), and the body weight was monitored every two days. The peripheral blood samples were obtained at predesignated timepoints, centrifuged at 1,000 × g for 10 min to extract plasma for quantification of serum cytokines, including IL-12 and IP-10, by ELISA kits (Peprotech). At day 8 post-administration, mice were sacrificed and the spleens were excised and weighted.

**NK cell isolation, culture, and activation.** NK cells were enriched from naïve mouse spleens with mouse NK cell isolation kit per the manufacturer's instructions. NK cells were cultured for 3 days in RPMI-1640 medium supplemented with 10% FBS in the presence of 20 ng $mL^{-1}$ mouse recombinant IL-2 and 50 ng $mL^{-1}$ IL-15. Then the NK cells were treated with $PEPA_{EE}$ or $SPEN_{EE}$-stressed and CT26-OVA-derived pyroptosomes for 24 h. Cells were harvested, and immuostained with APC anti-mouse NKp46 and PE anti-mouse CD107a antibodies for flow cytometry. The activated NK cells were further added into CT26-OVA cells at a ratio of 1:1, and incubated for 24 h. The cell killing efficacy of NK cells was evaluated by LDH assay.

### Supplementary Tables

**Table S1.** The characterization of synthetic polymer-IMDQ conjugates.

| polymer-IMDQ conjugates | Polymer-IMDQ | Polymer-TK-IMDQ |
|---|---|---|
| Repeating units of EPA[a)] | 48 | 49 |
| Repeating units of DPA[a)] | 47 | 46 |
| Repeating units of IMDQ[a)] | 7 | 7 |
| IMDQ loading contents [b)] | 9.56% | 9.89% |
| $M_n$ (kDa) [c)] | 23.45 | 23.50 |
| PDI [c)] | 1.38 | 1.41 |
| PPa conjugation [b)] | 7.26% | 7.74% |

[a)] Repeating units of methacrylate monomers was measured by $^1$H-NMR.

[b)] Drug loading capacity was measured by UV-Vis spectrum.

[c)] $M_n$ and PDI of the polymers were characterized by gel permeation chromatography (GPC).

### Supplementary Figures

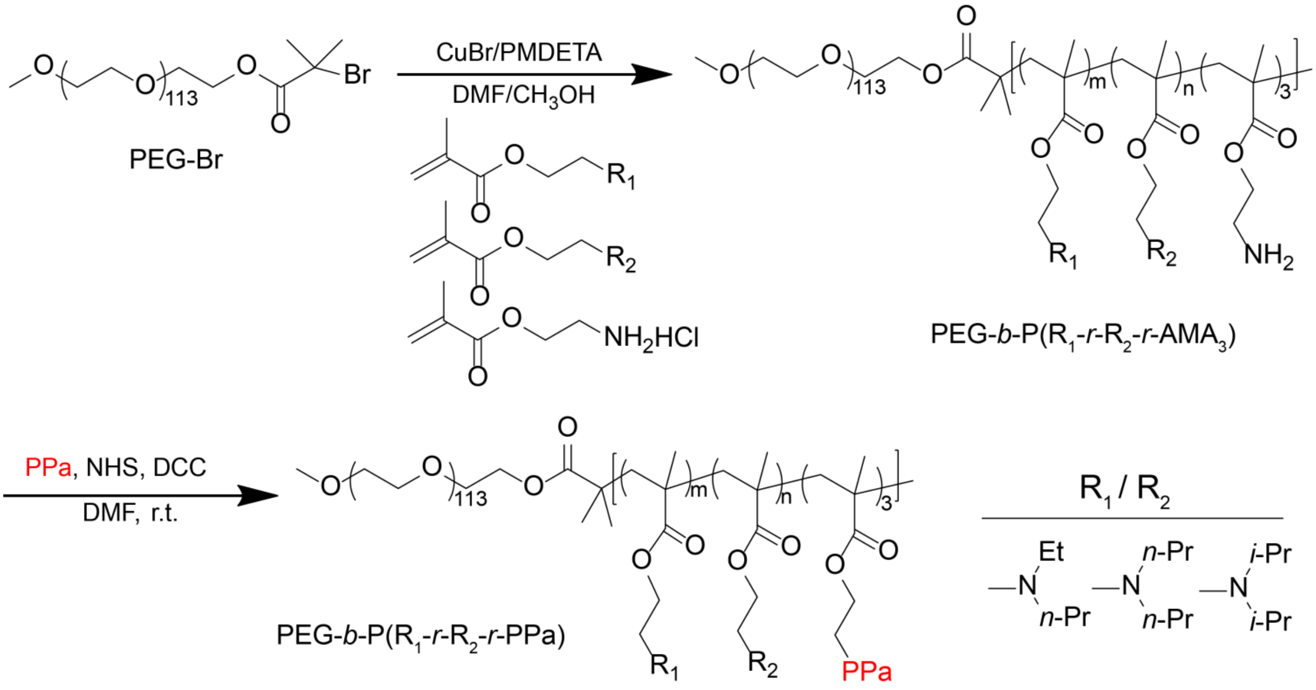


**Supplementary Fig. 1. Syntheses of ultra-pH-sensitive copolymers and PPa-conjugated polymers.**

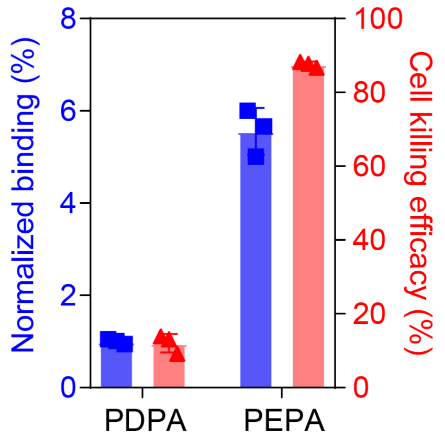


**Supplementary Fig. 2. Cell membrane binding (normalized to PDPA) and cell killing efficacy of PPa-conjugated PDPA and PEPA nanophotosensitizers at pH 6.0 on A549 cells.**

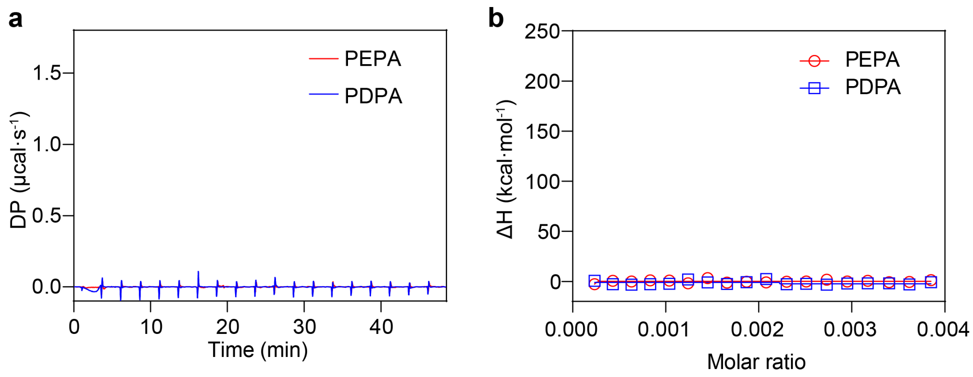


**Supplementary Fig. 3. The bio-nano interaction between the nanophotosensitizers and artificial lipid membrane measured by ITC at pH 7.4.**

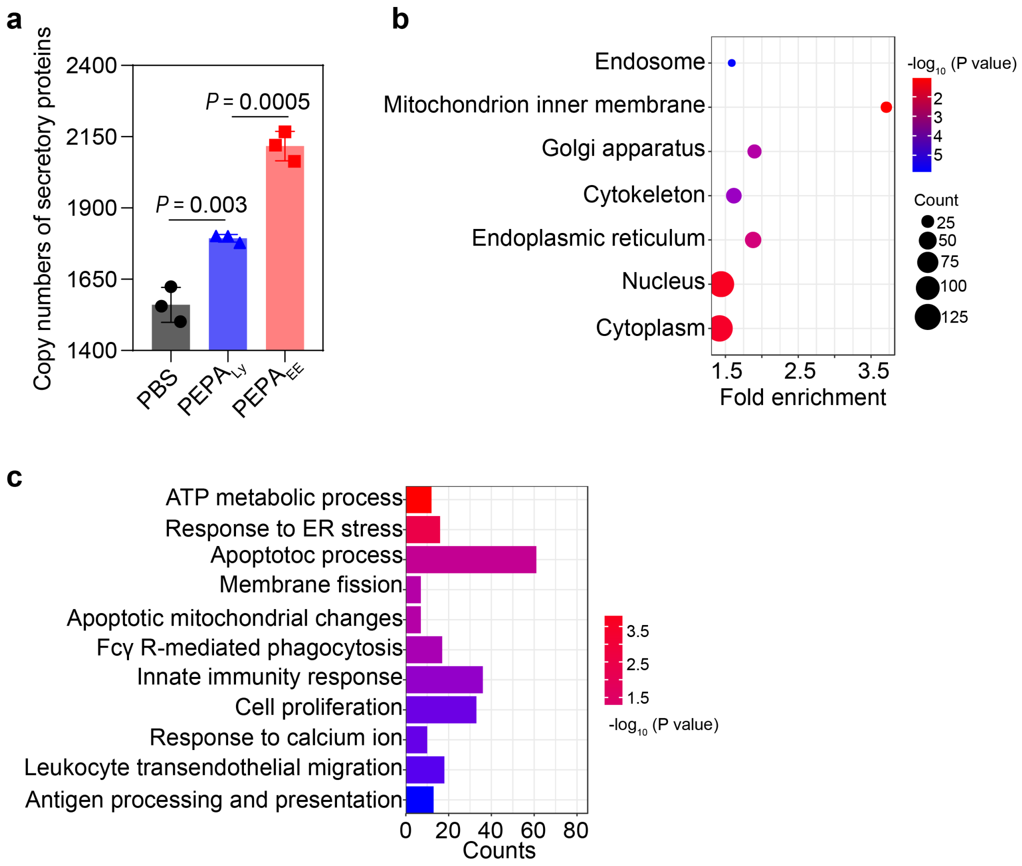


**Supplementary Fig. 4. Proteomics analysis of protein levels released from CT26 cells after treatment with PEPA-mediated EE or Ly stress. (a)** Copy number of secretory protein types. Data are shown as mean ± s.d. $n$ = 3 biologically independent experiment. Statistical significance was analyzed by one-way ANOVA followed by Tukey multiple comparisons test. **(b)** Enrichment of cellular component GOs specifically evoked by $PEPA_{6.5}$-mediated EE stress. **(c)** Clustering of significantly altered protein types between $PEPA_{EE}$ and $PEPA_{Ly}$.

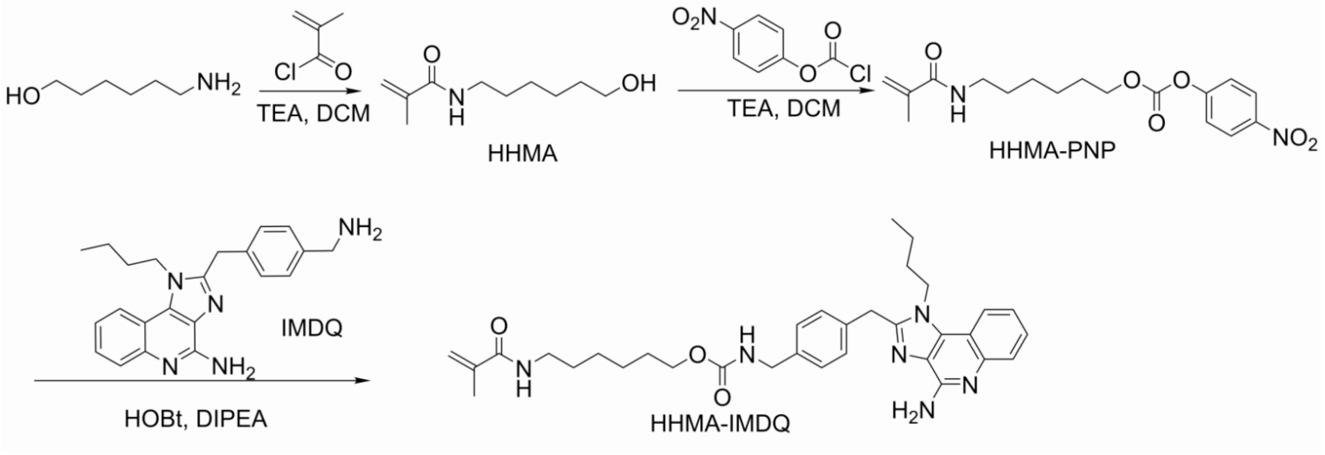


**Supplementary Fig. 5. Syntheses of the ROS-inactivated monomer HHMA-IMDQ.**

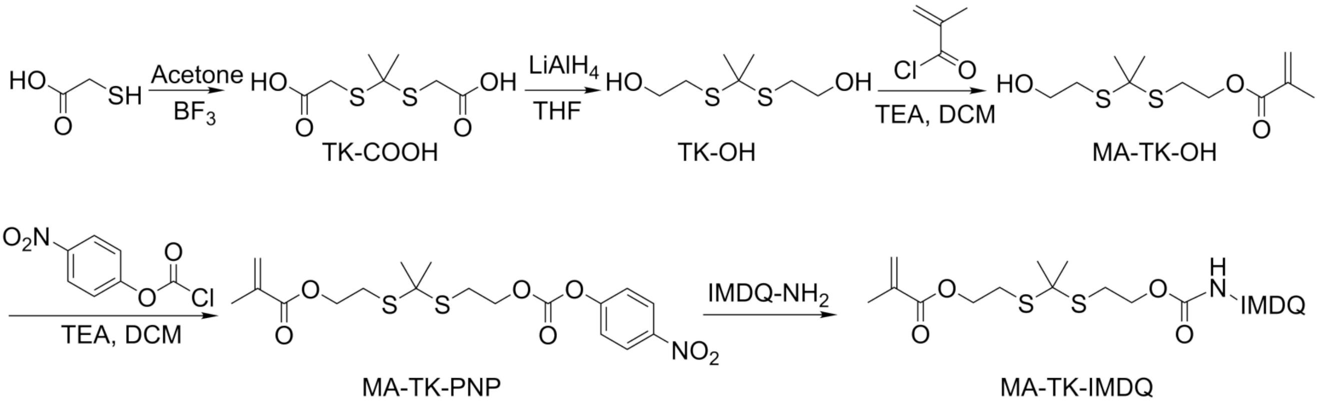


**Supplementary Fig. 6. Syntheses of the ROS-activatable monomer MA-TK-IMDQ.**

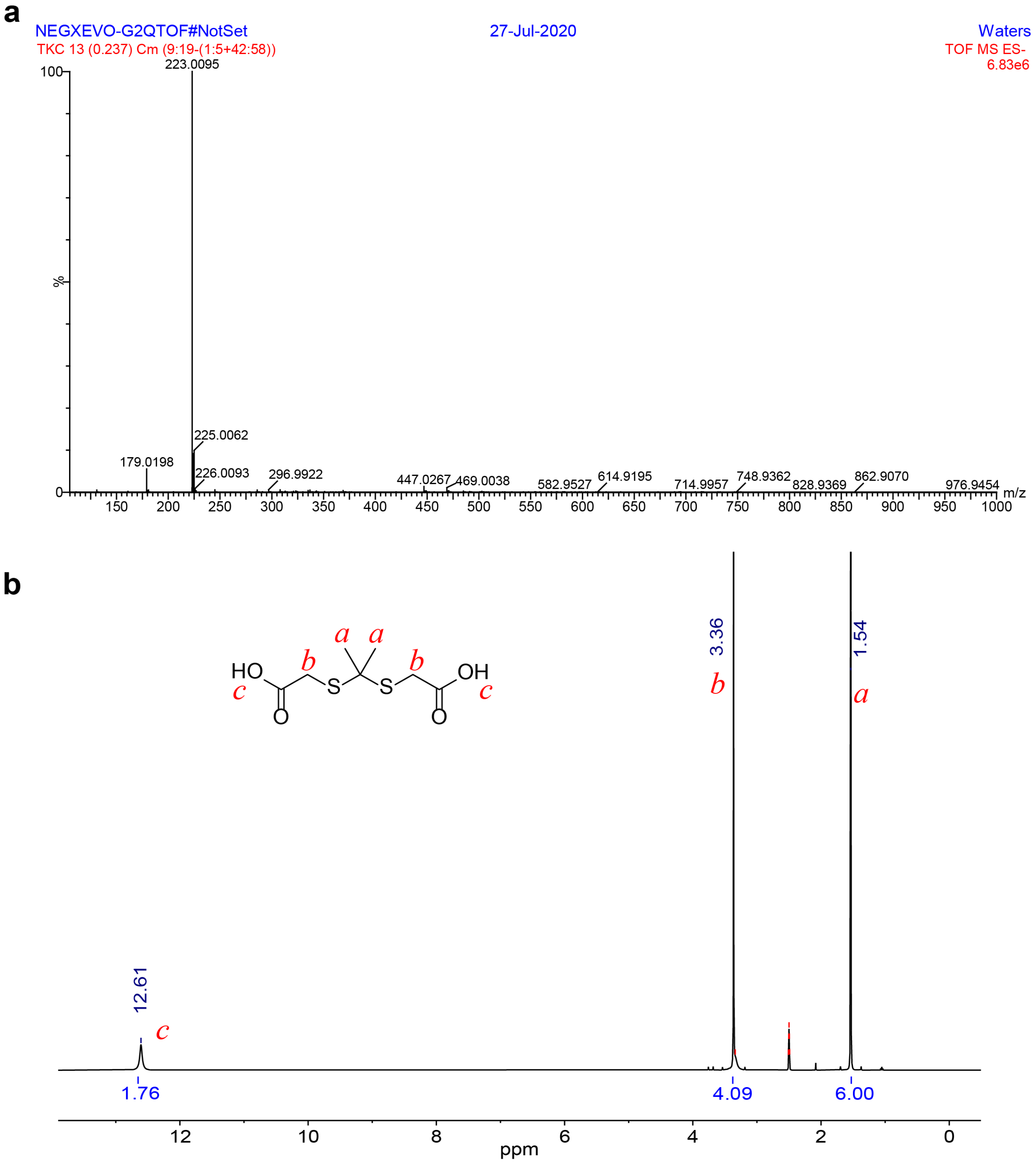


**Supplementary Fig. 7. Characterization of TK-COOH. (a)** ESI-MS spectra, m/z: 225 [M]$^+$, 223 [M-2H]$^+$. **(b)** $^1$H NMR spectra ($^1$H NMR (400 MHz, DMSO-$d_6$, δ): δ 12.61 (s, 2H), 3.36 (s, 4H), 1.54 (s, 6H).

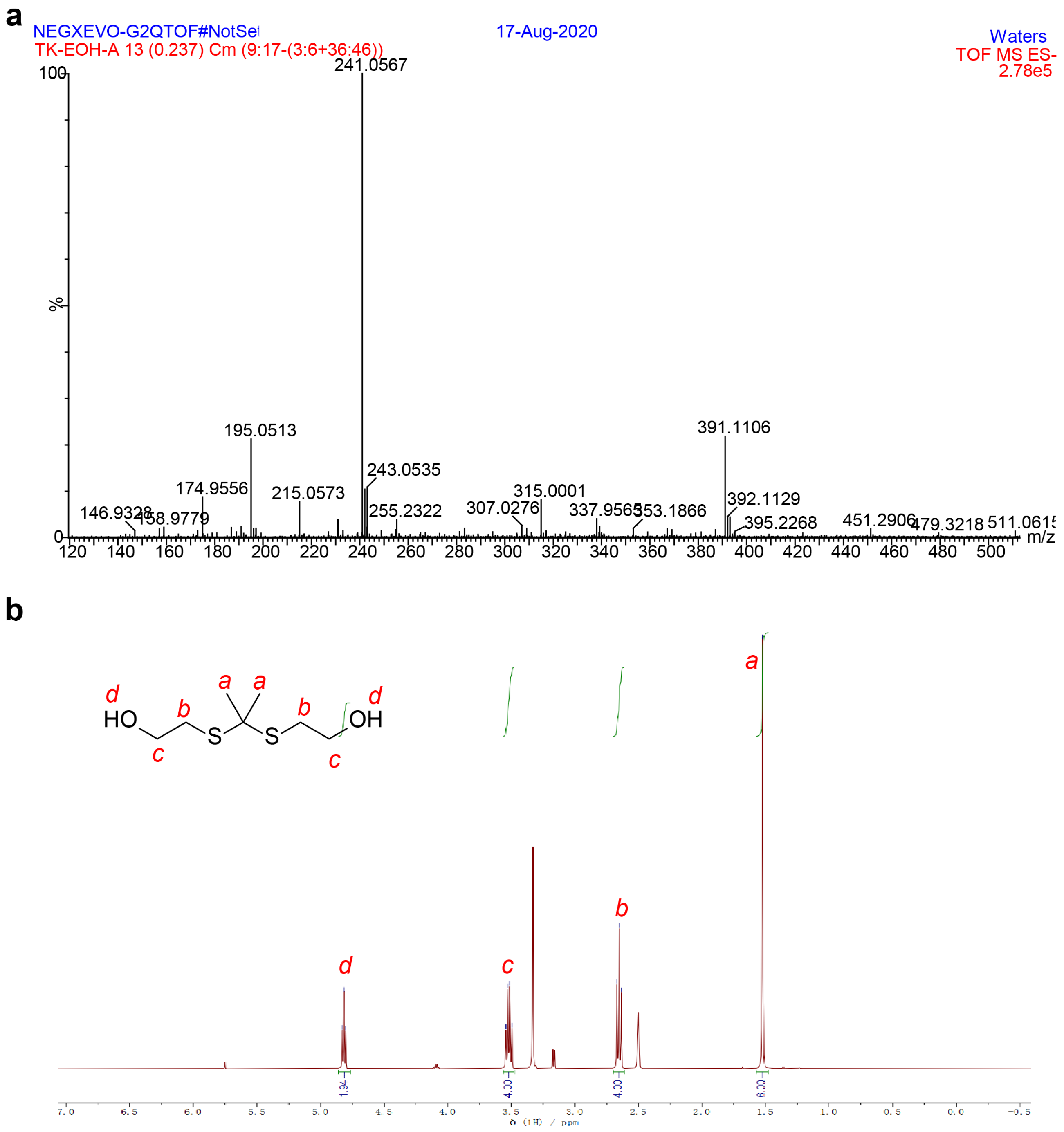


**Supplementary Fig. 8. Characterization of TK-OH. (a)** ESI-MS spectra, m/z: 195.0513 [M-H]$^-$, 241.0567 [M+HCOO]$^-$. **(b)** $^1$H NMR spectra ($^1$H NMR (400 MHz, DMSO-$d_6$, δ): 4.82 (t, 2H), 3.52 (q, 4H), 2.65 (t, 4H), 1.52 (s, 6H).

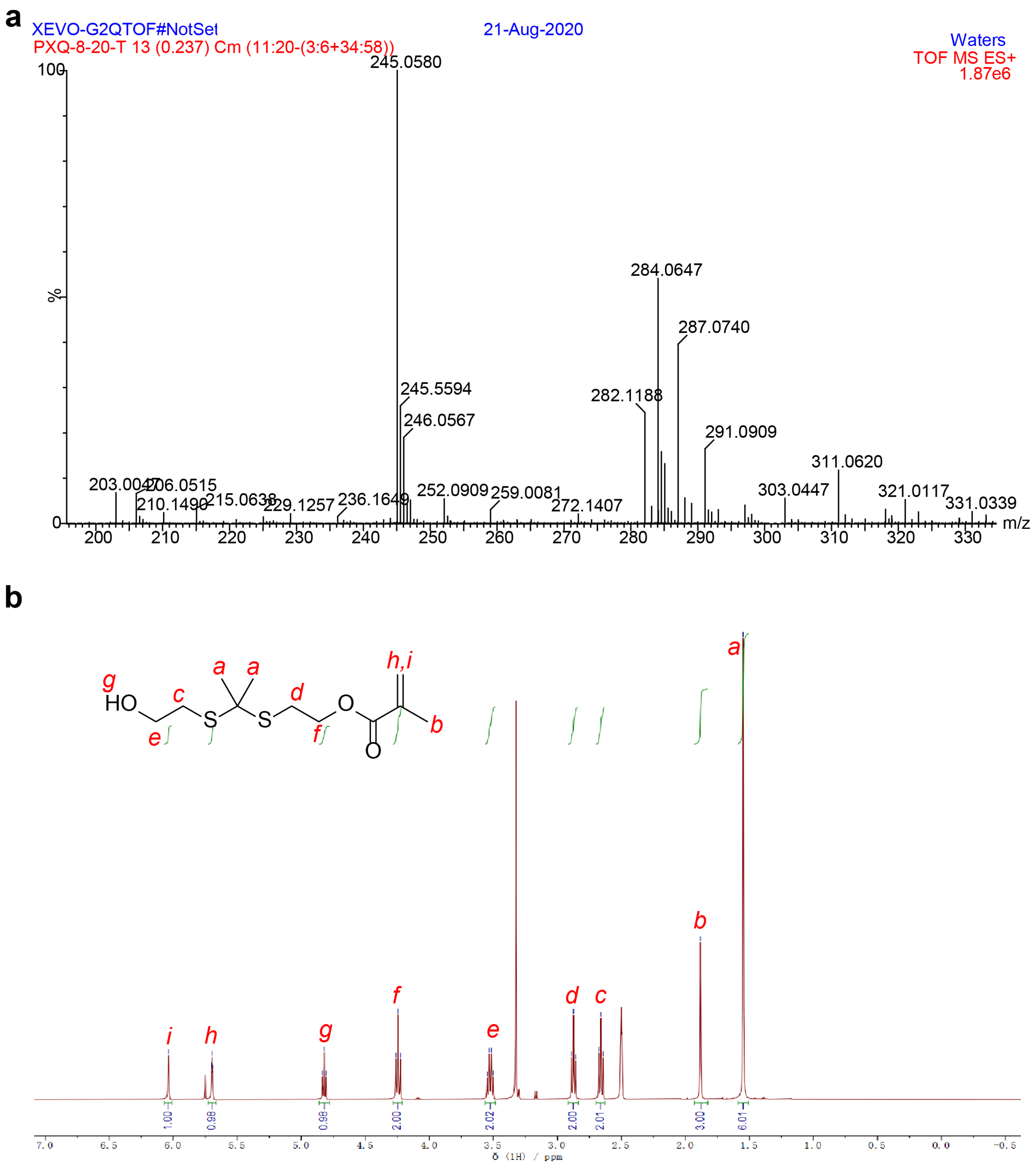


**Supplementary Fig. 9. Characterization of MA-TK-OH. (a)** ESI-MS spectra, m/z: 287.0740 $[M+Na]^+$. **(b)** $^1H$ NMR spectra ($^1H$ NMR (400 MHz, DMSO-$d_6$, δ): 6.04 (s, 1H), 5.70 (t, 1H), 4.82 (t, 1H), 4.24 (t, 2H), 3.52 (q, 2H), 2.87 (t, 2H), 2.66 (t, 2H), 1.88 (s, 3H), 1.55 (s, 6H).

**a**

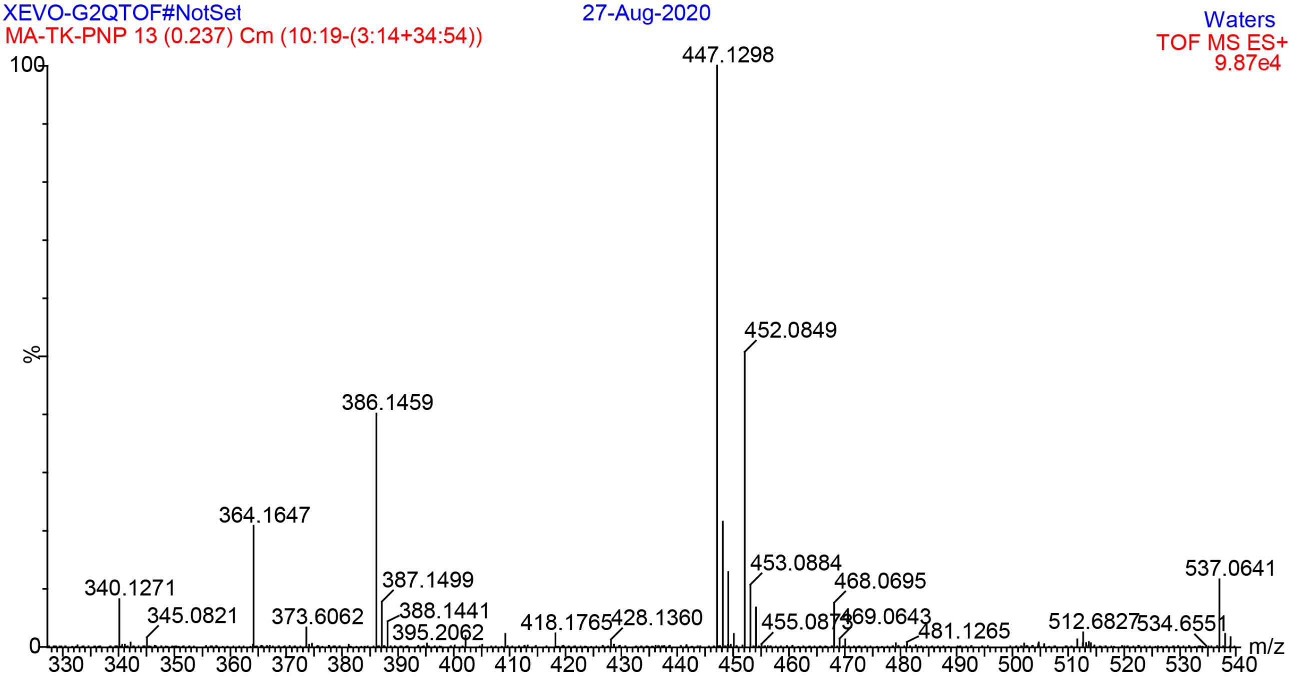


**b**

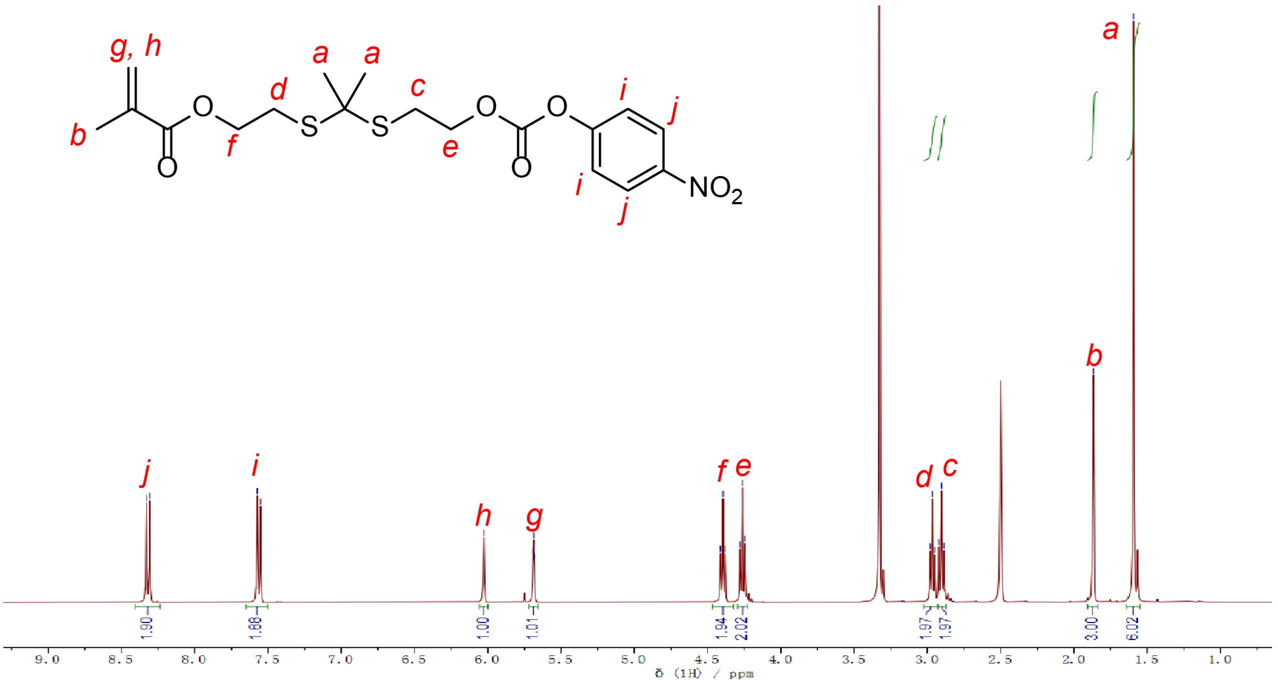


**Supplementary Fig. 10. Characterization of MA-TK-PNP. (a)** ESI-MS spectra, m/z: 447.1298 $[M+NH_4]^+$. **(b)** $^1H$ NMR spectra ($^1H$ NMR (400 MHz, DMSO-$d_6$, δ): 8.32 (d, 2H), 7.56 (d, 2H), 6.03 (s, 1H), 5.69 (t, 1H), 4.39 (t, 2H), 4.26 (t, 2H), 2.97 (t, 2H), 2.90 (t, 2H), 1.87 (s, 3H), 1.59 (s, 6H).

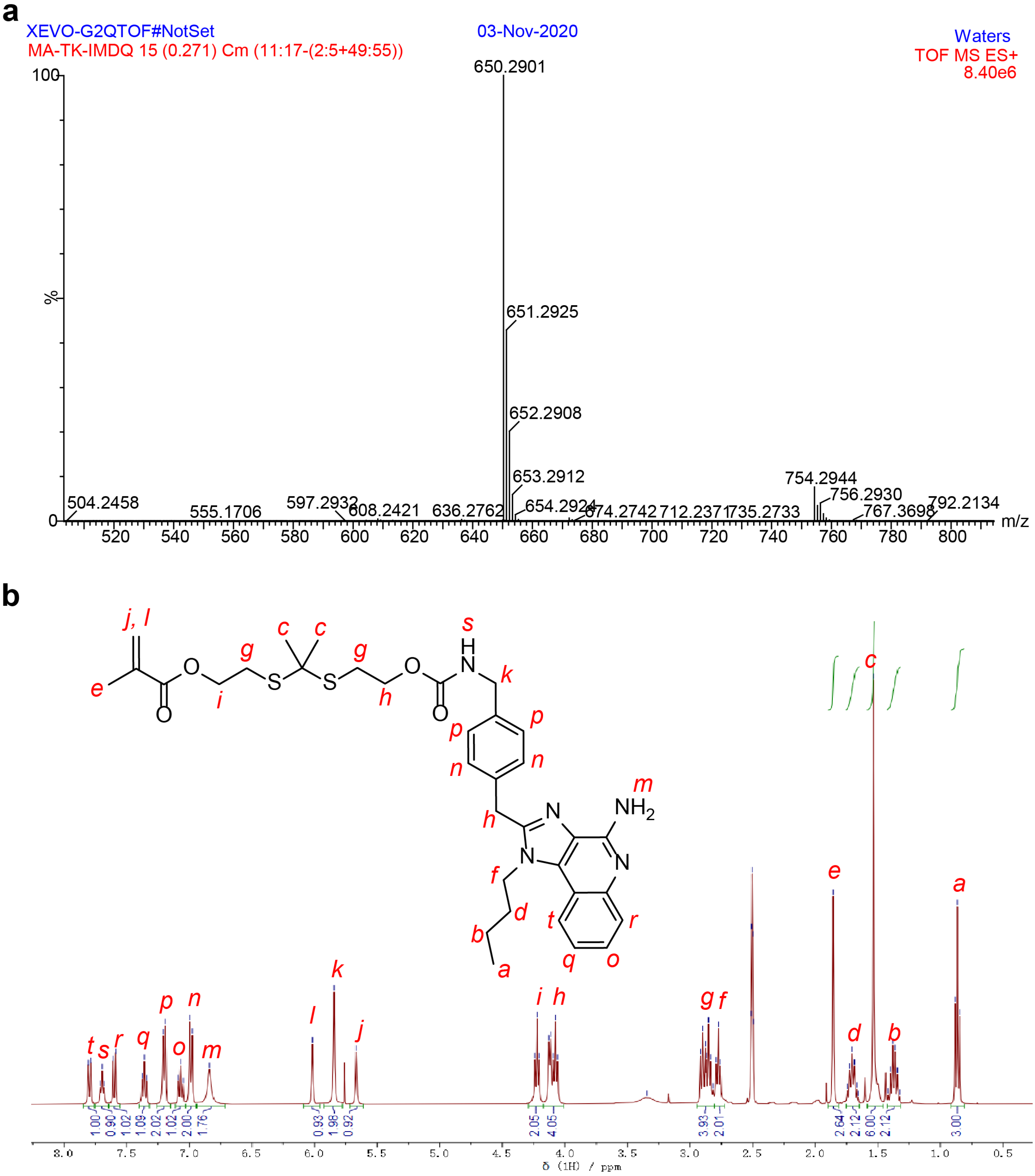


**Supplementary Fig. 11. Characterization of MA-TK-IMDQ. (a)** ESI-MS spectra, m/z: 650.2901 $[M+H]^+$. **(b)** $^1H$ NMR spectra ($^1H$ NMR (400 MHz, DMSO-$d_6$, δ): 7.80 (d, 1H), 7.70 (t, 1H), 7.60 (d, 1H), 7.36 (t, 1H), 7.20 (d, 2H), 7.07 (t, 1H), 6.99 (d, 2H), 6.84 (s, 2H), 6.02 (s, 1H), 5.84 (s, 2H), 5.67 (s, 1H), 4.22 (t, 2H), 4.09 (m, 4H), 2.86 (m, 4H), 2.76 (t, 2H), 1.86 (s, 3H), 1.70 (m, 2H), 1.53 (s, 6H), 1.37 (m, 2H), 0.86 (t, 3H).

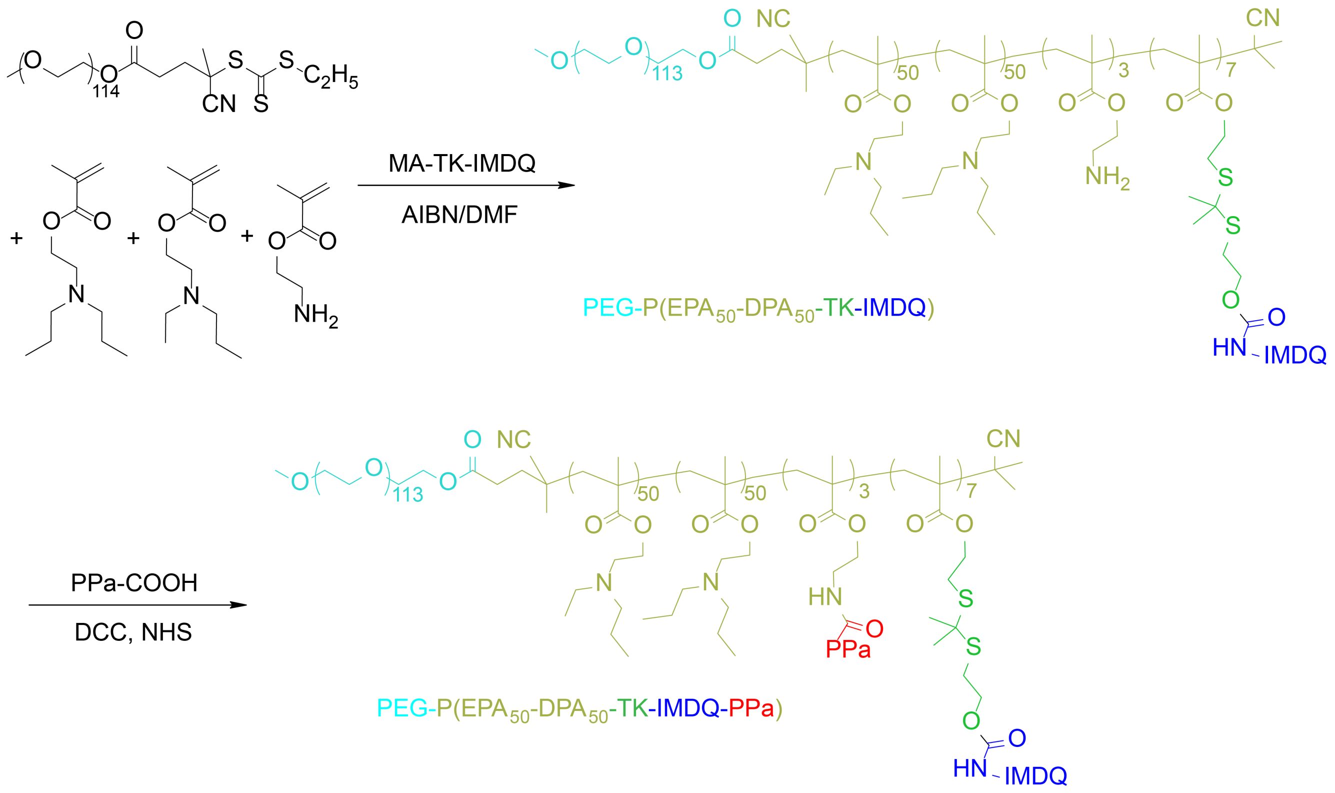


**Supplementary Fig. 12. Synthesis of polymer-IMDQ conjugates.**

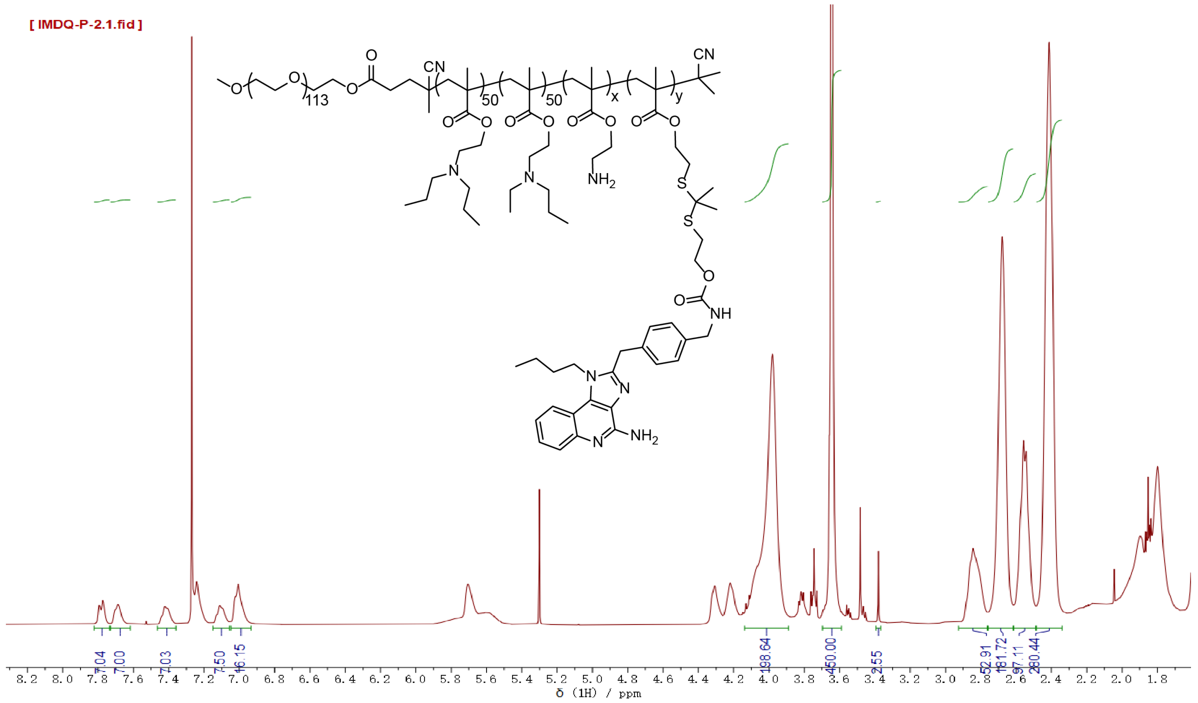


**Supplementary Fig. 13. $^{1}$H NMR spectra of PEG-*b*-P(EPA$_{50}$-*r*-DPA$_{50}$-*r*-TK-IMDQ-AMA$_{3}$).** $^{1}$H NMR, 400 MHz, DMSO-$d_6$. The peaks at 2.4 ppm and 2.5 ppm were used to estimate the EPA and DPA monomer compositions in the hydrophobic PR blocks. The peaks at 7.8 ppm, 7.7 ppm, and 7.4 ppm are used to quantify the conjugation efficacy of IMDQ monomer.

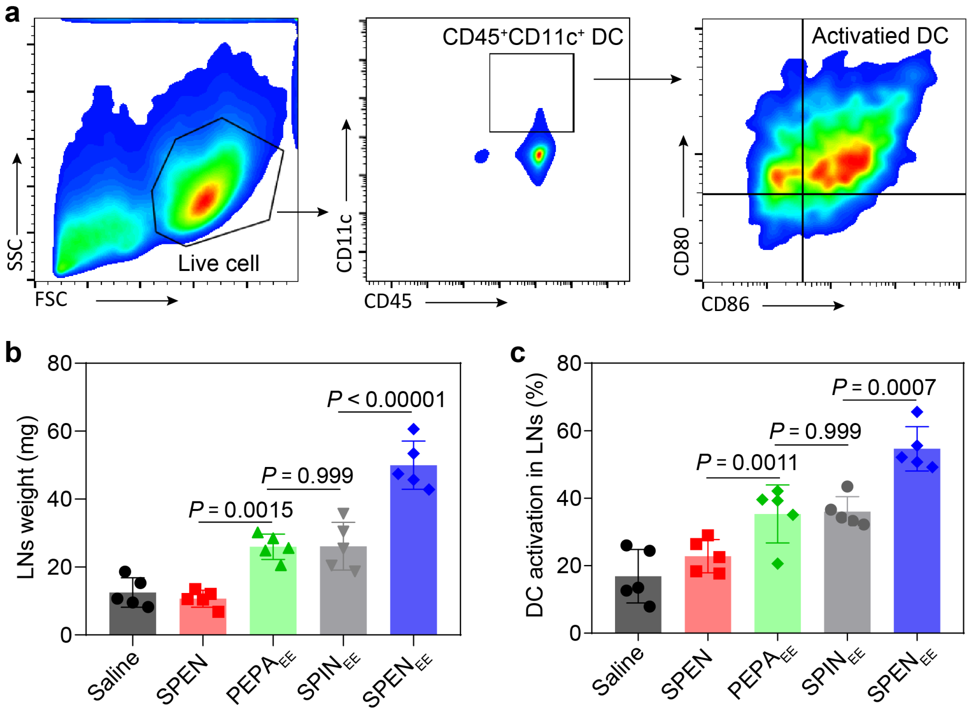


**Supplementary Fig. 14. In vivo DC activation in lymph nodes of mice immunized with pyroptosome. (a)** FACS gating strategy for matured DC. **(b)** Weight of the excised lymph nodes. **(c)** Quantification of the $CD80^{+}CD86^{+}$ DC in LNs. Data are shown as mean ± s.d.; *n* = 5 biologically independent mice; one-way ANOVA followed by Tukey multiple comparisons test.

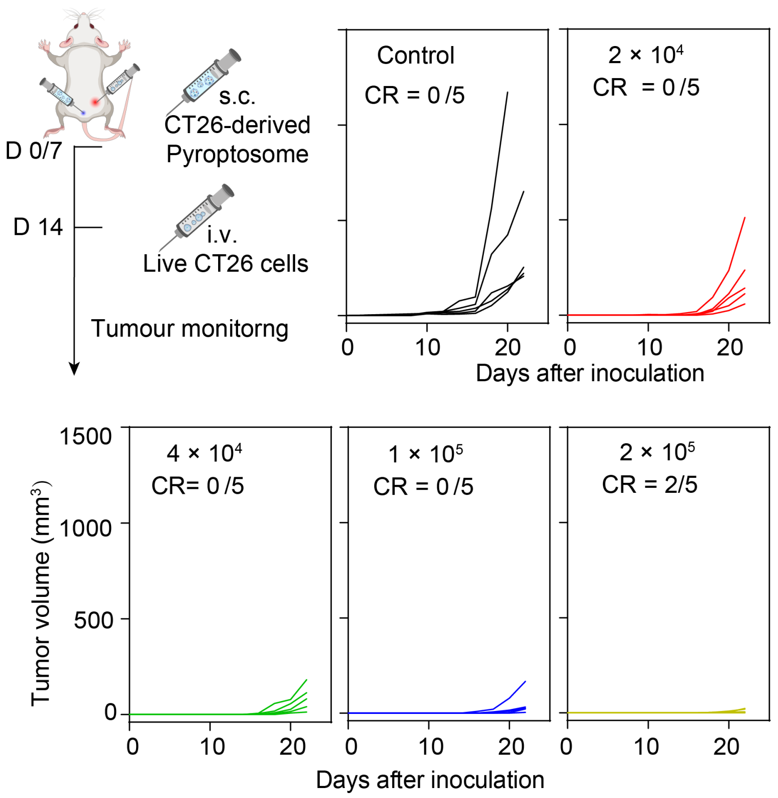


**Supplementary Fig. 15. In vivo vaccination capacity of pyroptosomes derived from SPEN$_{EE}$-evoked pyroptotic CT26 cells.** Individual tumour growth curves of CT26-tumour-bearing mice immunized with pyroptosomes. $n$ = 5 biologically independent mice.

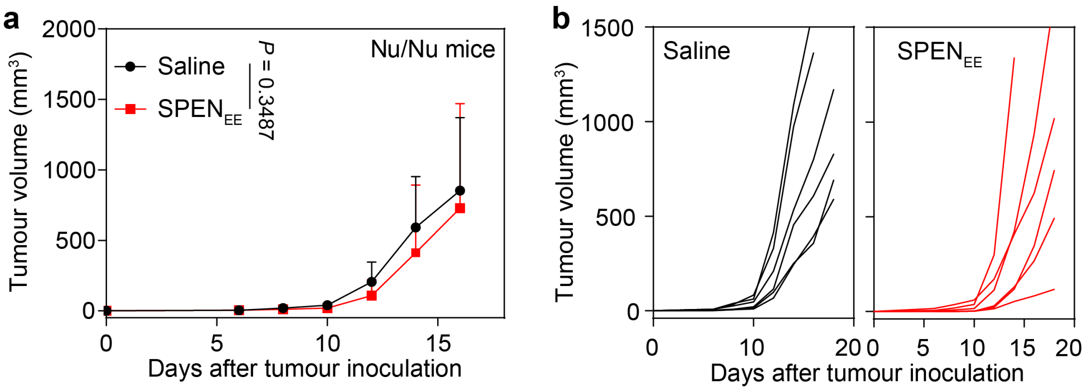


**Supplementary Fig.16. In vivo vaccination of pyroptosomes on immunodeficient Nu/Nu nude mice. Mice were immunized with pyroptosomes derived from SPEN$_{EE}$-evoked pyroptotic CT26 cells, and then inoculated with live CT26 cells. (a)** Average tumour growth curves. **(b)** Individual tumour growth curves. $n$ = 6 biologically independent mice; Two-way ANOVA followed by Bonferroni's multiple comparisons test

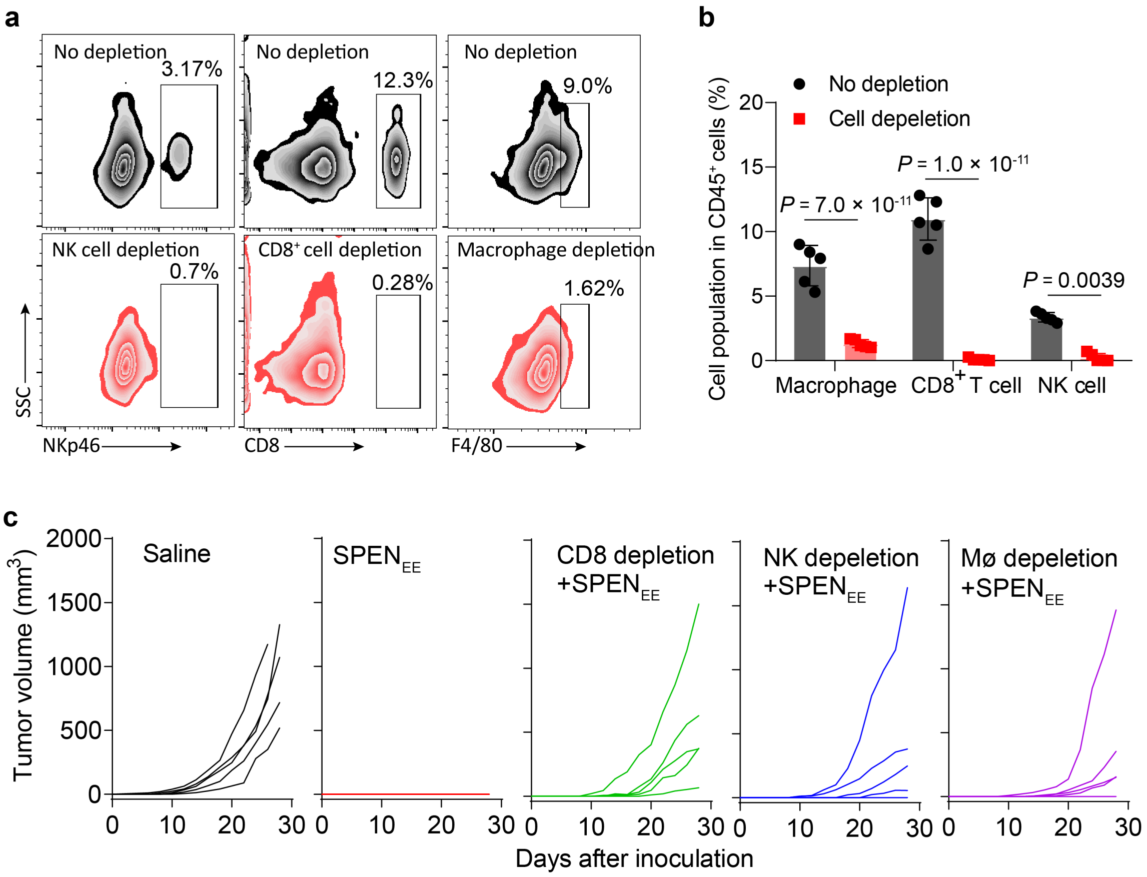


**Supplementary Fig. 17. Tumour inhibition of SPEN-mediated vaccination in CT26 tumour-bearing mice after depletion of CD8$^+$ T cells, NK cells, or macrophages, respectively. (a)** Representative flow cytometry plots and **(b)** quantification of NK cells, CD8$^+$ T cells, and macrophages in peripheral blood of mice after depletion of specific immune cells. **(c)** Individual tumour growth curves. Data are shown as mean ± s.d. *n* = 5 biologically independent mice.

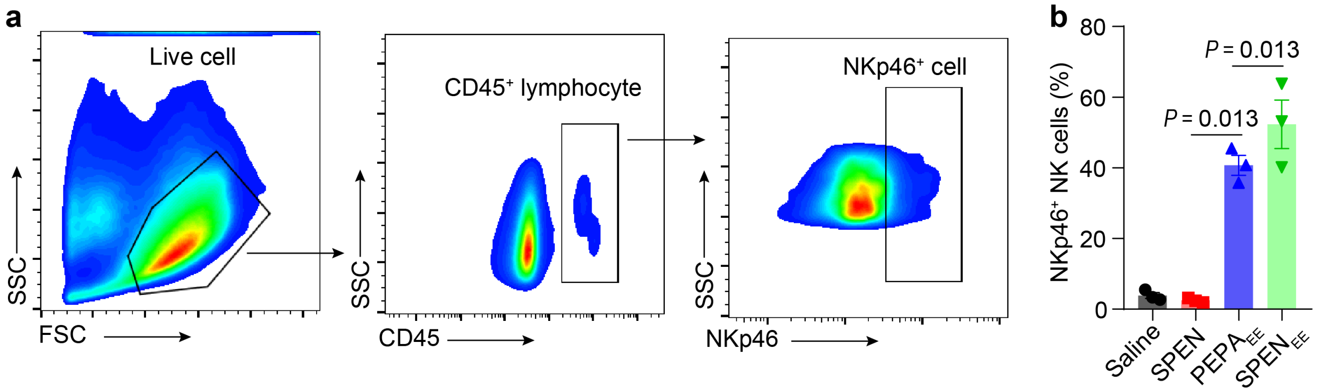


**Supplementary Fig. 18. In vitro NK activation with the treatment of pyroptosome. (a)** FACS gating strategy for activated NK. **(b)** Quantification of NKp46$^+$ cells among isolated CD45$^+$ lymphocyte. Data are shown as mean ± s.d.; *n* = 3 biologically independent experiment.

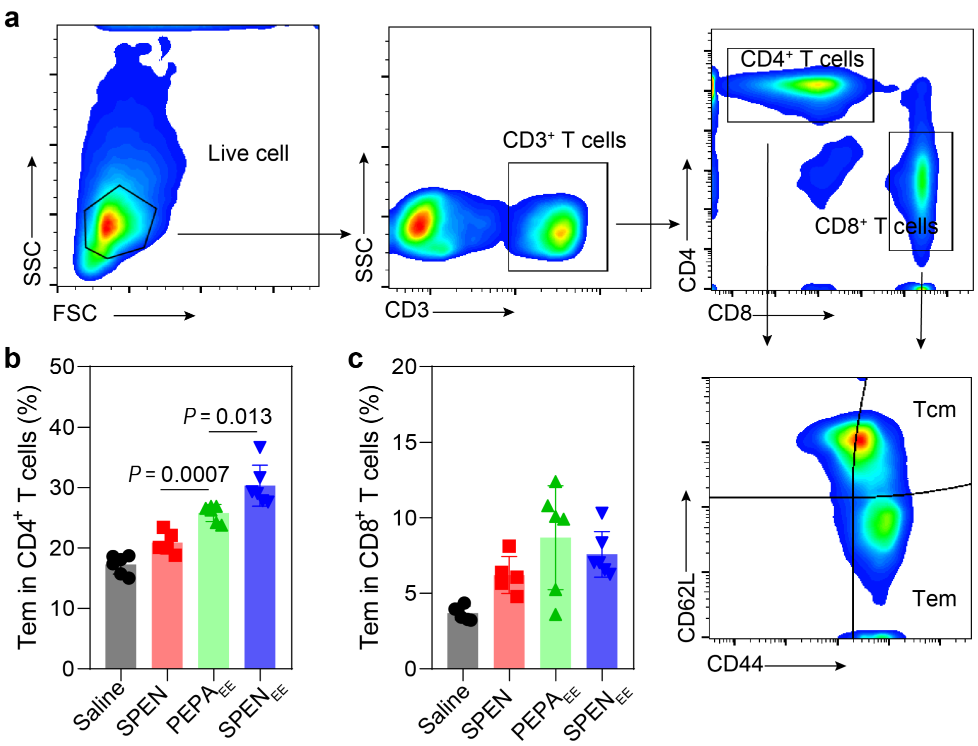


**Supplementary Fig. 19. Quantification of effector memory T cells (Tem) in MC38 tumour-bearing mice immunized with SPEN-mediated in situ vaccination. (a)** FACS gating strategy for memory T cells. **(b)** Quantification of Tem in $CD4^+$ T cells. **(c)** Quantification of Tem in $CD8^+$ T cells. $n = 6$ biologically independent mice; one-way ANOVA followed by Tukey multiple comparisons test.

### Supplementary Videos

**Supplementary video 1**. Rapid membrane ballooning and pyroptosis triggered by PEPA-mediated EE stress on CT26-GFP cells. CT26-GFP cells were treated with PEPA (1 µg/mL PPa) for 0.5 h, and immediately irradiated with a 660 nm laser (6 J $cm^{-2}$). Shown is the video of representative field recorded immediately after irradiation (The exact time duration, h: min: s: ms).
